# The microevolutionary response to male-limited X-chromosome evolution in *Drosophila melanogaster* reflects macroevolutionary patterns

**DOI:** 10.1101/800672

**Authors:** Jessica K. Abbott, Adam K. Chippindale, Edward H. Morrow

**Affiliations:** Department of Biology, Section for Evolutionary Ecology, Lund University, Sölvegatan 37, 22362 Lund, Sweden; Biology Department, Queen’s University, Kingston, Ontario, Canada, K7L 3N6; School of Life Sciences, University of Sussex, John Maynard Smith Building, Brighton, UK, BN1 9QG

**Keywords:** sexual conflict, microarray, gene expression, experimental evolution

## Abstract

Due to its hemizygous inheritance and role in sex determination, the X chromosome is expected to play an important role in the evolution of sexual dimorphism, and to be enriched for sexually antagonistic genetic variation. By forcing the X chromosome to only be expressed in males over >40 generations, we changed the selection pressures on the X to become similar to those experienced by the Y. This releases the X from any constraints arising from selection in females, and should lead to specialization for male fitness, which could occur either via direct effects of X-linked loci or trans-regulation of autosomal loci by the X. We found evidence of masculinization via upregulation of male-benefit sexually antagonistic genes, and downregulation of X-linked female benefit genes. Interestingly, we could detect evidence of microevolutionary changes consistent with previously documented macroevolutionary patterns, such as changes in expression consistent with previously established patterns of sexual dimorphism, an increase in the expression of metabolic genes related to mitonuclear conflict, and evidence that dosage compensation effects can be rapidly altered. These results confirm the importance of the X in the evolution of sexual dimorphism and as a source for sexually antagonistic genetic variation, and demonstrate that experimental evolution can be a fruitful method for testing theories of sex chromosome evolution.

## Introduction

Sex chromosomes have long been of interest to researchers in evolution and genetics, due to their special mode of inheritance and association with sex determination (Abbott et al. 2017). In XY systems, the X chromosome is expected to be feminized (enriched in female-biased genes compared to the autosomes) and/or demasculinized (impoverished for male-biased genes compared to the autosomes) because it spends two thirds of its time in females, compared to only half for autosomes. Although there are exceptions, these predictions generally seem to hold true for species with heteromorphic sex chromosomes (reviewed in Dean and Mank 2014). The X is also expected to be enriched for sexually antagonistic genetic variation, i.e. alleles with opposite effects on fitness between males and females (also known as intralocus sexual conflict). However expectations from theories of sexual antagonism are slightly different, in that they predict that the X should be enriched for loci with recessive male-benefit and/or dominant female-benefit sexually antagonistic alleles (Rice 1984, Patten 2018; but see Fry 2009). This is because any sexually antagonistic male-benefit allele will be expressed in hemizygous males, but masked in heterozygous females when recessive. Although this prediction is more challenging to test, it has some support (Connallon and Knowles 2006, Dean et al. 2012).

These two phenomena are not mutually exclusive, and although these predictions (enrichment both of female-biased genes and of male-benefit loci on the X) may superficially appear to be conflicting, this is only the case if we assume that female-biased genes only benefit females, and male-biased genes only benefit males. This is not necessarily the case; e.g. a gene that suppresses a male-specific pathway should be upregulated in females, but this doesn’t imply that the pathway itself is female-benefit. Consistent with this, Immonen *et al.* (2017) found that female seed beetles upregulated expression of male-biased genes and downregulated expression of female-biased genes after mating. Another important factor is time scale. New male-biased or male-benefit genes probably arise at equal rates across the genome, but given that many new mutations are recessive, X-linked loci have an initial advantage in that they will immediately be expressed in males and can be positively selected (Charlesworth et al. 1987). However due to the fact that the X spends more time in females than in males, it may be a less favourable environment than the autosomes for male-benefit or male-biased genes (but see Patten 2018). This leads to selection for such genes to re-locate to the autosomes over evolutionary time, causing demasculinization of the X. Indeed, previous results have shown that young testis-specific genes tend to be located on the X more often than expected, but that old genes are less likely to be located on the X than expected (Zhang et al. 2010). Similarly, Long *et al.* (2012) reviewed traffic to and from the X, and found that genes often move from the X to the autosomes in both Diptera and mammals.

Hemizygous inheritance has other implications as well. Because males only have one copy of the X while females have two, males are expected to have half the expression at X-linked loci as females do, all else being equal. In many species this asymmetry in expression seems to be disadvantageous, since various forms of dosage compensation have evolved (Chandler 2017). For example, female mammals inactivate one X chromosome, after which both sexes hyperexpress the X to maintain equal expression with the autosomes (Graves 2016). In *Drosophila*, males instead hyperexpress the X directly, through the action of dosage compensation complexes that bind to high-affinity sites (Conrad and Akhtar 2012). The dosage compensation complex prevents chromatin formation, allowing more efficient hyperexpression of X-linked loci close to the high-affinity sites. This seemingly elegant system may however constitute a constraint for males, in that it hinders the evolution of sex-specific expression, particularly in loci close to high-affinity sites. This has been demonstrated in a cross-species comparison, which found that male-biased genes tended to be located in regions without dosage compensation, farther away from high-affinity sites (Bachtrog et al. 2010). Finally, theory suggests that mitonuclear interactions may be particularly likely on the X (Ågren et al. 2018), although empirical support for this prediction varies across species (Dean et al. 2014). Nevertheless, Rogell *et al.* (2014) found that mito-sensitive genes (i.e. nuclear genes that changed in expression when paired with different mitochondrial genotypes) are underrepresented on the *Drosophila* X, even if nuclear genes with a mitochondrial annotation (i.e. mito-nuclear genes) are not (Dean et al. 2014).

Experimental alteration of mating regimes has been successfully employed to test hypotheses related to sexual selection and sexual conflict and their effect on gene expression (Hollis et al. 2014, Immonen et al. 2014, Innocenti et al. 2014, Perry et al. 2016). Some of these studies support the idea that elevated levels of sexual selection/conflict result in a shift towards the male optimum via an increase in expression of male-biased genes (e.g. Hollis et al. 2014), but others do not (e.g. Immonen et al. 2014, Veltsos et al. 2017). In contrast, studies of sex chromosome evolution are often observational or comparative in nature (e.g. Mank and Ellegren 2009, Wright et al. 2015), due to the difficulty in constructing experimental tests of macroevolutionary patterns (although there are some exceptions; Dean et al. 2012). Manipulative experiments testing predictions about sex chromosome evolution are therefore particularly valuable, since they make it possible to disentangle causal effects from stochastic effects (Kawecki et al. 2012, Abbott et al. 2017). We carried out a male-limited X chromosome evolution experiment in *Drosophila melanogaster* designed to integrate predictions from both sexual antagonism and sex chromosome evolution, where X chromosomes were passed from father to son for >40 generations, and never expressed in females. This experimental protocol changes the selection pressures on the X to become similar to those on the Y chromosome, and should result in specialization of evolved X’s to enhance male fitness. It also decouples inheritance of the X and the mitochondria, which may have implications for mitonuclear conflict. We therefore expected to see masculinization at the phenotypic level, as well as in the expression profile via upregulation of male-benefit genes and downregulation of female-benefit genes. Based on previous evidence of sexual antagonism in this species, we also expected to see antagonistic changes in fitness in males and females (Prasad et al. 2007).

Overall, our results were consistent with these predictions, although there were some surprises. Evidence of masculinization and sexual antagonism were relatively weak on the phenotypic level but there were clear signatures of both on gene expression. We were also able to detect a change in locomotory activity consistent with previous characterization of this trait as sexually antagonistic (Long and Rice 2007). Interestingly, Gene Ontology analysis suggested changes in some traits which are not previously characterized as sexually antagonistic in this species but likely to be important in sexual selection, such as vision and learning/memory. An exciting finding, given the short timescale of the experiment, was evidence of microevolutionary changes consistent with the patterns of macroevolutionary change discussed above, including upregulation of male-biased genes and downregulation of female-biased genes, change in the expression of metabolic genes related to mitonuclear conflict, and evidence that dosage compensation may constitute a constraint for male-benefit genes. These results suggest that we were successful in releasing males from constraints arising from selection in females, and therefore experimentally manipulating the selection pressures on the X chromosome.

## Methods

### Experimental protocol

All populations were derived from the LH_M_ stock (Chippindale and Rice 2001), which is a large, outbreeding population with an easily replicable maintenance regime (see Supplementary Methods for more details). Fly lines were maintained so as to follow the ancestral culturing protocol as closely as possible. In order to control inheritance of X-chromosomes and limit expression of the X to males, we used compound X (i.e. “double X” or DX) females. DX females have two marked X-chromosomes that are linked at the centromere (*C(1)DX, y, f*), and were backcrossed to the LH_M_ stock population for 6 generations before the start of the experiment, so that they carried a random LH_M_ Y-chromosome and LH_M_ autosomes which were >98% identical to the base stock. When wildtype males are mated with DX females, X-bearing sperm can fertilize Y-bearing eggs, resulting in father-son transmission of the X (autosomal inheritance is unaffected; Figure 1). Six populations were started simultaneously; three replicate male-limited X-chromosome (MLX) populations and three control populations. All populations started by mating 480 males from the LH_M_ stock population to an equal number of females. In the control populations these females were wildtype LH_M_ females, and in the MLX populations they were the backcrossed DX females. The total initial population size was therefore 960 individuals, but this was later reduced to 640 individuals (320 of each sex) for logistical reasons (Abbott et al. 2013). The X-chromosome population size was inevitably lower in the MLX populations compared to the Control populations under this protocol, since females in the MLX populations do not carry wildtype X-chromosomes. However adjusting for this difference would have resulted in large differences in the amount of autosomal variation available between the treatments, so we instead chose initial population sizes deemed large enough to provide a reasonable level of X-linked standing genetic variation. Because the X is in a hemizygous state in males, we also simultaneously started a “recombination box” treatment for each replicate MLX population (Prasad et al. 2007, Abbott et al. 2013). The recombination box serves to introduce sufficient recombination to reduce the effects of linkage, such as background selection and hitchhiking. See the Supplementary Methods for more information.

**Figure 1:**
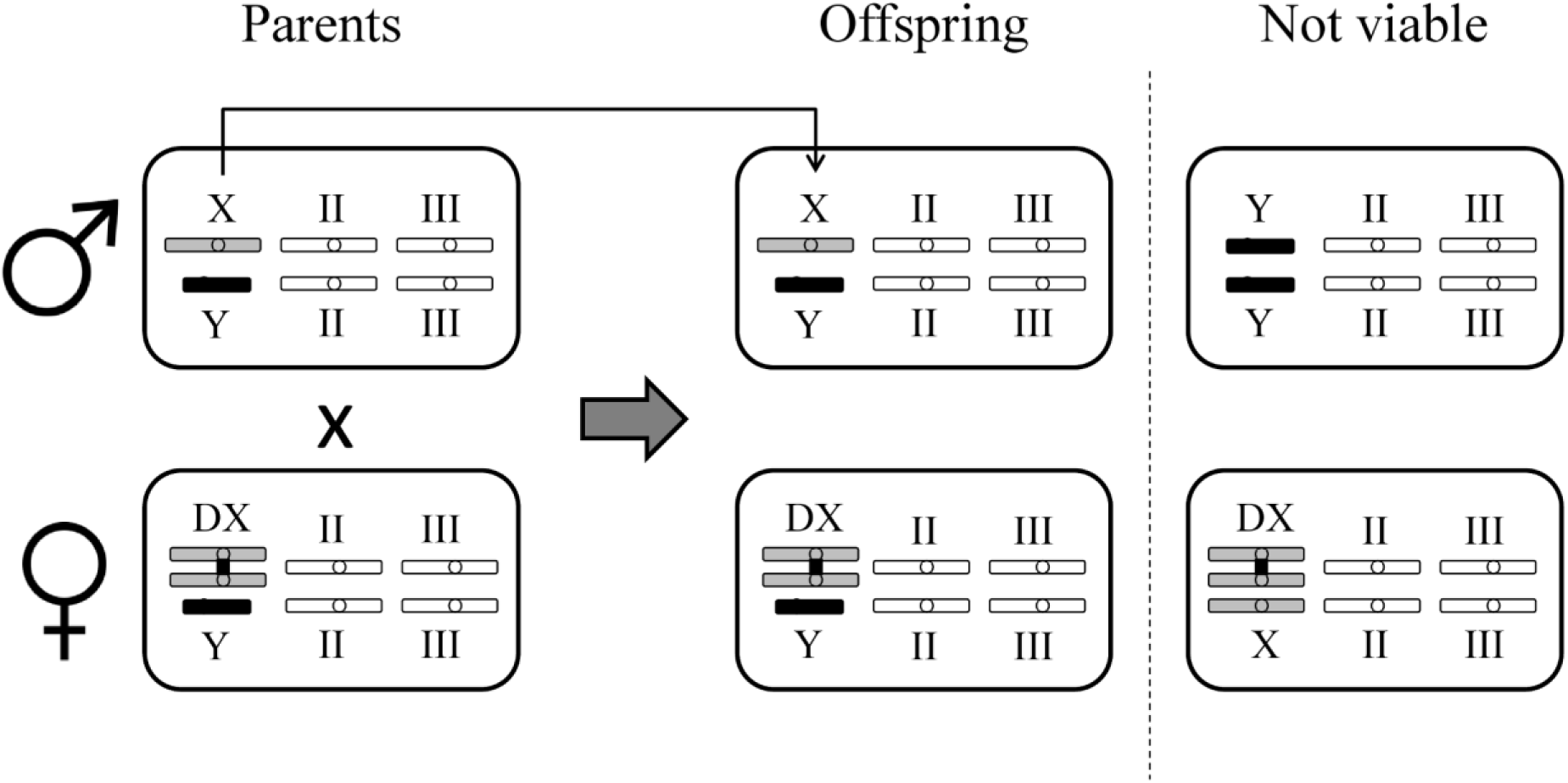
MLX evolution protocol. Males are mated to females with a double X-chromosome (DX), which forces father-son transmission of the X-chromosome, and produces wildtype males with a paternally inherited X-chromosome and a maternally inherited Y-chromosome. New DX females with a paternally inherited Y-chromosome are also produced. Triple-X and double-Y individuals are not viable.

### Phenotypic assays

After 40 generations of experimental evolution, we measured male fitness using a standard eye-marker protocol (Abbott et al. 2010, Abbott et al. 2013). See the Supplementary Methods for more details. Target males were taken directly from the MLX and Control populations, as were Control females. However wildtype females are not normally produced within the MLX populations, so they were obtained by first crossing MLX males with females from the Control population of the same number. This results in females with one MLX X-chromosome and one Control X-chromosome. Although the observed effects of the MLX selection on females might have been larger if using females that have two MLX X-chromosomes, this also admits the possibility of (likely deleterious) inbreeding effects on the X. We therefore elected to use females that were heterozygous with respect to X-chromosome origin, in order to avoid confounding inbreeding effects with sexual antagonism.

We were also interested to see if MLX evolution had effects on sex-specific survival from egg to adult, and therefore also collected data on offspring sex ratios when measuring fitness. For female data, test-tubes were kept after eggs were counted and total number of adult offspring was later recorded in order to provide some data on total survival rates. Finally, based on results from the gene expression analysis, indicating that locomotory activity might be of interest (see below), we also collected data on that trait across sexes and selection treatments. We used a protocol from a previous study of locomotory activity in the LH_M_ population (Long and Rice 2007). In the set-up, flies were kept in mixed-sex groups in which the target sex is wildtype and the opposite sex is brown-eyed. Density was kept at the same level as in the adult competition vials, 16 individuals of each sex. In order to avoid observer bias (e.g. that the eye is drawn to motion), vial volume was divided up into 6 sections and the section to be observed was determined using a random number generator. The first target fly detected in the section was then observed for 8 seconds and scored for locomotory behaviour (binary 0/1 score if the behavior was observed or not). If no target fly was found in the selected section then the observation was immediately terminated and a new random number was generated. Observations were repeated until all vials have been observed 10 times, and an average score per vial (i.e. proportion active flies) was used for further analysis.

In the statistical analysis of the phenotypic data, we preferred to carry out the analysis on population means since the population is the appropriate level of replication for this experiment. However results from an alternative approach using raw data with population as a random factor nested within treatment were qualitatively similar in most cases, and are reported in the supplementary information. Means were calculated from data for 20 vials per sex, selection treatment, and replicate population for fitness, sex ratio, and survival. Since vials means are already based on ten repeated observations in the locomotory activity assay, only 5 vials per sex, selection treatment, and replicate population were used. Fitness data were mean-zero unit-variance standardized to allow comparison between the sexes, and analysed with a two-way ANOVA with sex and selection treatment as fixed factors. Because sex ratio and locomotory activity data are in the form of proportions, these were analysed using GLMs with a quasibinomial distribution and sex and selection treatment as fixed factors. Similarly, survival was analysed with a GLM with a quasibinomial distribution, but because survival was only measured in offspring from the female part of the fitness assay, selection treatment was the only factor in this analysis. Posthoc analysis of interaction effects was carried out using the “lsmeans” package (Lenth 2016). All analyses were carried out in R (Team 2014).

### Gene expression analysis

Gene expression data was collected after 50 generations of experimental evolution using microarrays. RNA was extracted with Trizol (Invitrogen) and purified using an RNeasy Mini Kit (Qiagen). On day 12 from egg, two replicates of 8 individuals were taken for each combination of sex, selection treatment, and replicate population. RNA quantity and quality was assessed using an Agilent Bioanalyzer (Agilent Technologies) prior to sample preparation and hybridisation at the Uppsala Array Platform (using GeneChip Drosophila Genome 2.0 Affymetrix microarrays following the manufacturer’s instructions). Note that the male samples in this analysis partially overlap with data published elsewhere (Abbott et al. 2013), but the datasets were normalized and analysed separately and results suggest that these two sets of analyses can be considered independent in terms of their output (see Supplementary Results).

Analysis was carried out in BioConductor (http://www.bioconductor.org), and data were pre-processed using Robust Multichip Average (RMA) in the “affy” package (Gautier et al. 2004). Significant differences in gene expression levels were tested using a model that included selection treatment, sex, and their interaction, as well as replicate population to control for non-independence of replicate samples, with a false discovery rate (FDR) correction at 0.05. Replicate population was considered a random factor nested within treatment. In line with other studies of expression changes after experimental evolution (Immonen et al. 2014, Veltsos et al. 2017), we did not employ a fold-change cutoff since most changes were expected to be relatively small.

Once genes that had changed significantly in expression between selection treatments had been identified, they were then divided into downregulated and upregulated sets and these sets were further investigated separately. In cases where gene lists were very short (e.g. genes that showed a significant interaction effect), properties were examined manually using information from online databases from the National Center for Biotechnology Information (NCBI; https://www.ncbi.nlm.nih.gov/). Upregulated and downregulated genes were tested for Gene Ontology (GO) terms, chromosomal distribution, tissue-specificity, and association with sex-specific fitness and sexual antagonism, as measured in a previous study of the LHm population (Innocenti and Morrow 2010). Overrepresentation of GO categories was analysed using hypergeometric testing (“hyperGTest” in R), which tests whether particular GO terms are associated with a gene set more often than expected by chance. Chromosomal distribution was tested using a χ^2^ test (“chisq.test” in R). Chromosomal location tests were run both with and without including chromosome 4, since this chromosome is non-recombining and may skew results of the χ^2^ analysis. Tissue specificity was measured in the same manner as in previous expression analyses of the LH_M_ population (Innocenti and Morrow 2010, Abbott et al. 2013), with Bonferroni correction for multiple testing. Association with sex-specific fitness and sexual antagonism was analysed using two-tailed mean-rank gene set enrichment (MR-GSE) tests (Innocenti and Morrow 2010, Abbott et al. 2013).

### Relationship to extant sexual dimorphism

We tested if change as a result of the selection treatment was in the same direction as existing expression differences between the sexes (i.e. whether MLX selection had a masculinizing effect, such that male-biased genes became even more upregulated and female-biased genes became even more downregulated). For this, we ran a model of the effect of sex on expression in the Control samples only, then compared direction of change in the selected populations with direction of sexual dimorphism in the Control population using a χ^2^ test. Although these data could potentially also be analyzed with a linear model, distributions of the datapoints are bimodal due to exclusion of non-significant expression differences (see Figure 2), making a χ^2^ test more appropriate. We checked whether associations with sex-bias were robust to source population differences by testing if sex-biased genes (as reported in the Sebida database, Gnad and Parsch 2006) and sexually discordant loci (Stocks et al. 2015) were overrepresented among genes that responded to the selection treatment using χ^2^ tests (Collet et al. 2016). We also tested for enrichment of genes more recently reported as sexually antagonistic based on alleleic variation rather than expression differences (Ruzicka et al. 2019) in this dataset, as well as for enrichment of genes that have previously been shown to change expression in response to manipulations of the intensity of sexual conflict (Immonen et al. 2014, Innocenti et al. 2014). Enrichment was again tested using χ^2^ tests.

**Figure 2:**
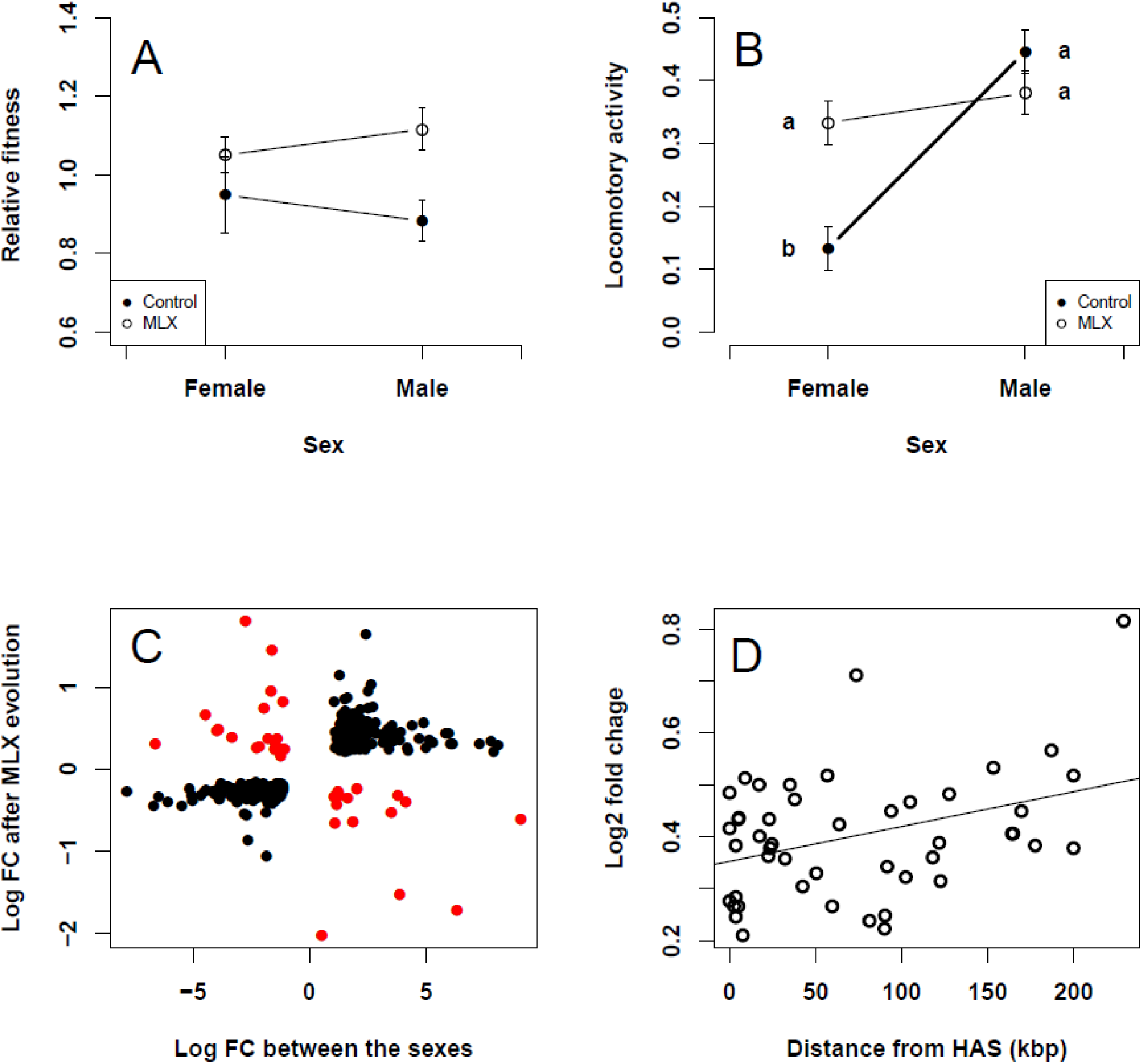
Overview of main results. A: Fitness in relation to selection treatment and sex. MLX males have higher fitness than Control males, but the treatments overlap in females. B: Comparison of direction of sexual dimorphism in expression in Control populations, with change in selected populations. Positive values on the x-axis indicate male-biased expression, and positive values on the y-axis indicate upregulation after MLX evolution. Changes as a result of MLX evolution are much more often in the same direction as extant sexual dimorphism (black) compared to being in opposite directions (red). C: Locomotory activity in relation to experimental evolution treatment and sex. Homogenous subsets are indicated with letters. MLX females have activity levels that are not significantly different from male activity levels. D: Relationship between change in expression after MLX evolution, and distance from the closest high-affinity site (HAS). High-affinity sites are associated with dosage compensation, and genes that were located farther away from high-affinity sites appear to be less constrained in their response to the selection treatment.

Male-biased genes are more likely to be located outside dosage compensation regions (Bachtrog et al. 2010), so we hypothesized that there might be a relationship between distance to high-affinity sites (associated with dosage compensation) and change in expression level as a result of the selection treatment using linear models. Location of high-affinity sites was obtained from Straub et al. (2008). Visual inspection of the relationship between change in expression and distance to the nearest high-affinity site revealed a potential outlier in the upregulated dataset, and this was confirmed using a standard diagnostic (Cook’s distance > 1). Results excluding this outlier are therefore reported here, but the relationship is still highly significant if it is included.

### Mitonuclear conflict and interaction analyses

We also checked if any of the transcripts identified as having changed as a result of male-limited selection showed enrichment of genes previously identified as potentially subject to mitonuclear conflict using χ^2^ tests (Rogell et al. 2014). Finally, two interaction network analyses were carried out, firstly to investigate degree of overlap with a previous dataset, and secondly to determine whether most significant genes fall into several large or many small clusters. Details of these interaction analyses are included in the Supplementary Information.

## Results

### Phenotypic assays

Fitness was standardized by sex before analysis, so there was no effect of sex on fitness (as expected) and no significant interaction between sex and selection treatment. However there was a main effect of selection treatment on fitness (*P* = 0.039, Table S1), such that MLX populations had higher fitness than Control populations. Although the interaction term was not significant, the difference between the treatments was larger for males than for females (Figure 2A).

There was a significant effect of the interaction between sex and selection treatment on offspring sex ratio (*P* = 0.0009), as well as significant main effects (Table S2). Sex ratio was female-biased overall, but the MLX populations produce a higher proportion male offspring than Control populations. This difference is especially noticeable for offspring of females with an MLX X-chromosome (Figure S2). A slightly female-biased sex ratio is not unusual for the LH_M_ population (Long and Pischedda 2005), but the effect was likely compounded by overcrowding that occurred as a result of higher-than-anticipated female fecundity for this assay. In addition, the lower values overall for females with an MLX X-chromosome are a result of the fact that, for logistical reasons, these vials were counted two days later than the male vials. Some additional male mortality appears to have occurred during this period, which made us unable to distinguish the eye colour in the offspring and unambiguously assign male offspring to targets or competitors. The magnitude of the advantage of an MLX X-chromosome to male survival is therefore probably an overestimate relative to normal culturing conditions.

There was a significant effect of selection treatment on offspring survival (*P* = 5.41*10^−05^), where offspring of MLX females had higher survival than offspring of control females (Table S2, Figure S3). Based on the results from the sex ratio analysis, it seems likely that this difference is a result of increased survival in sons who inherit an MLX X-chromosome, assuming that the sex ratio at fertilization is the same across selection treatments.

There was a significant effect of the interaction between sex and selection treatment on adult locomotory activity (*P* = 7.23*10^−05^), as well as a significant main effect of sex (Table S2). Males had higher locomotory activity than females overall, but this activity was increased in MLX females, such that they were as active as males (Figure 2C).

Results from the alternative approach using raw data with population as a random factor nested within treatment were qualitatively similar in all cases, except for fitness (Table S3).

### Changes in gene expression

6286 transcripts were variable in expression across samples and were retained for analysis. Of these, 6276 showed a significant effect of sex, and 518 a significant effect of selection treatment. There was only one transcript showing a significant selection effect, that did not also show a significant sex effect (CG14957, annotated as being related to chitin production https://www.ncbi.nlm.nih.gov/gene/38386). This suggests that most of the variation across samples is related to sexual dimorphism. Only 4 transcripts showed a significant interaction effect between sex and selection treatment: CG10514 gene product from transcript CG10514-RA (CG10514), Glutaminyl cyclase (QC), PCI domain-containing protein 2 (PCID2), and sallimus (sls). Because there were so few interaction effects, only transcripts showing an effect of the selection treatment are considered further. We report results for upregulated and downregulated transcripts separately (i.e. increased or decreased in MLX lines compared to Control), since they proved to be qualitatively different.

### Up-versus downregulated transcripts

Of the 518 transcripts that responded significantly to the selection treatment, 342 were upregulated and 176 were downregulated. Log_2_ fold changes ranged from ~0.15 to approximately 2.

Both upregulated and downregulated transcripts were distributed non-randomly across chromosomes. Upregulated genes were slightly enriched on the X but considerably enriched on chromosome 4 (χ^2^ = 94.9102, df = 3, *P* < 2.2e-16, Table 1). This significant result is apparently driven by the enrichment on chromosome 4, since the difference is no longer significant when chromosome 4 is excluded from the analysis (χ^2^ = 0.7212, df = 2, *P* = 0.6973). Downregulated genes were underrepresented on chromosome 2 but enriched on the X-chromosome, consistent with the expectation that female-benefit genes might be overrepresented on the X (Table 1). This pattern showed a trend towards significance when all data was included (χ^2^ = 6.4141, df = 3, *P* = 0.09311), and became marginally significant when chromosome 4 was excluded (χ^2^ = 6.408, df = 2, *P* = 0.0406).

**Table 1:** Chromosomal location of MLX transcripts. Up-regulated transcripts were located on chromosome 4 significantly more often than expected (highlighted). Because chromosome 4 does not recombine, this is likely due to genetic hitchhiking. Down-regulated transcripts were located on the X chromosome more often than expected (highlighted). This is consistent with the expectation that dominant female-benefit loci should often be located on the X.

| Chromosome | Up-regulated transcripts |  | Down-regulated transcripts |  |
| --- | --- | --- | --- | --- |
|  | Observed | Expected | Observed | Expected |
| X | 47 | 54.7 | **40** | **28.2** |
| 2 | 130 | 133.5 | 59 | 68.7 |
| 3 | 148 | 150 | 75 | 77.2 |
| 4 | **16** | **2.1** | 1 | 1.1 |

Upregulated genes had tissue-specific expression (as previously characterized in Chintapalli et al. 2007) more often than expected in the testes, ovaries, virgin and mated spermathecae, hindgut, fat body, heart, and salivary glands (Table S4). Down-regulated genes showed tissue-specific expression in the crop only (Table S4).

GO terms for upregulated and downregulated genes are reported in Tables S5 and S6 respectively. Many of these were rather general and uninformative (e.g. “System process”, “Signaling”, “Secretion”). Some interesting exceptions for upregulated genes are terms associated with locomotory behaviour, metabolism, larval behaviour, phototaxis, learning and memory, and various terms related to vision (response to light stimulus, photoreceptor differentiation, detection of visible light). For downregulated genes, there were a number of terms associated with DNA replication and damage repair, cell cycle regulation, and oogenesis.

We tested whether genes that responded to the selection treatment were non-randomly associated with fitness, as measured in a previous study (Innocenti and Morrow 2010). We found that upregulated genes were significantly associated with increased male fitness, decreased female fitness, and sexual antagonism (Table 2). Downregulated genes were significantly associated with increased female fitness and decreased male fitness, but there was no overrepresentation of genes characterized as sexually antagonistic (i.e. simultaneously increasing female fitness and decreasing male fitness, or vice versa; Table 2).

**Table 2:** Relationship with fitness. MLX transcripts show a pattern of association with fitness that is remarkably consistent with theory. Up-regulated transcripts are good for male fitness, bad for female fitness, and significantly sexually antagonistic. Down-regulated transcripts are good for female fitness and bad for male fitness. Fitness data obtained from Innocenti & Morrow (2010).

| Genes associated with | Up-regulated transcripts | Down-regulated transcripts |
| --- | --- | --- |
| Male fitness | Positive ($P = 1.9299 \times 10^{-22}$) | Negative ($P = 0.0099$) |
| Female fitness | Negative ($P = 8.0925 \times 10^{-07}$) | Positive ($P = 6.8031 \times 10^{-07}$) |
| Sexual antagonism | Positive ($P = 2.2462 \times 10^{-17}$) | No association ($P = 0.3947$) |

### Relationship to extant sexual dimorphism

We found that changes in expression were overwhelmingly consistent with the direction of extant sexual dimorphism (483/518 in the same direction, versus 35/518 in the opposite direction, χ^2^ = 387.5, df = 1, *P* = 2.2*10^−16^; Figure 2B). Similarly, sex-biased genes (both male-biased and female-biased as classified in SEBIDA) were overrepresented among the genes changed as a result of the selection treatment. Upregulated genes were male-biased more often than expected (χ^2^ = 237.8, df = 2, *P* = 2.2*10^−16^), and downregulated genes were female-biased more often than expected (χ^2^ = 193.1, df = 2, *P* = 2.2*10^−16^; Table 3). There was no evidence of enrichment of genes recently characterized as sexually antagonistic (all *P* > 0.07) by Ruzicka et al (2019). Genes that have been previously shown to respond to altered intensity of sexual conflict in this species (Innocenti et al. 2014) were not overrepresented in this dataset, and were if anything underrepresented among downregulated genes (χ^2^ = 4.8698, df = 1, *P* = 0.02733). Genes that responded to altered intensity of sexual conflict in *Drosophila pseudoobscura* (Immonen et al. 2014) were somewhat overrepresented in this dataset (100 genes found where 74 were expected; χ^2^ = 10.199, df = 1, *P* = 0.001405).

**Table 3:** Direction of change in expression after MLX evolution in relation to sex bias in gene expression. Male-biased genes are significantly overrepresented among up-regulated transcripts, and female-biased genes are significantly overrepresented among down-regulated transcripts.

| Sex bias | Up-regulated transcripts |  | Down-regulated transcripts |  |
| --- | --- | --- | --- | --- |
|  | Observed | Expected | Observed | Expected |
| Male | **176** | **69.5** | 13 | 47.2 |
| Unbiased | 49 | 89.1 | 4 | 55.3 |
| Female | 15 | 81.4 | **146** | **60.5** |

Interestingly, we found a significant effect of distance to high-affinity sites (HAS) on expression of upregulated genes. Increased distance from HAS regions resulted in greater upregulation as a result of the selection treatment (*F*_*1,45*_ = 7.2502, *P* = 0.0099; Figure 2D). There was no overlap with genes previously identified as highly sexually concordant or discordant in their effect (Stocks et al. 2015).

### Mitonuclear conflict

We examined three classes of genes potentially subject to mitonuclear conflict, after Rogell *et al.* (2014): mito-annotated genes (Gene Ontology ID 0005739), mito-sensitive genes (Innocenti et al. 2011), and mito-proteome genes (Lotz et al. 2014). Although there was no signature of overrepresentation of these classes among the genes that responded to the selection treatment as a whole (for mito-sensitive genes: χ^2^ = 2.179, df = 1, *P* = 0.14; for mito-proteome genes: χ^2^ = 0.076, df = 1, *P* = 0.78), and mito-annotated genes were in fact slightly underrepresented (χ^2^ = 4.108, df = 1, *P* = 0.043), the pattern of regulation was revealed to be skewed. Both mito-annotated genes and mito-sensitive genes were upregulated as a result of the selection protocol more often than expected by chance (mito-annotated: χ^2^ = 7.629, df = 1, *P* = 0.0057; mito-sensitive: χ^2^ = 20.02, df = 1, *P* = 7.65*10^−6^; table 4). In addition, an analysis of chromosome location that was carried out for upregulated mito-annotated genes (the only class of gene that had sufficient sample size for such an analysis), revealed that there was significant overrepresentation of genes located on chromosome 4 (6/44; χ^2^ = 58.95, df = 3, *P* = 9.88*10^−13^).

**Table 4:** Direction of change in expression after MLX evolution for three classes of mitonuclear genes. Mito-sensitive and mito-annotated genes were upregulated significantly more often than expected. There was no deviation from the null expectation for mito-proteome genes.

| Regulation | Mito-sensitive |  | Mito-annotated |  | Mito-proteome |  |
| --- | --- | --- | --- | --- | --- | --- |
|  | Observed | Expected | Observed | Expected | Observed | Expected |
| Upregulated | **44** | **29.3** | **6** | **2.6** | 11 | 13.7 |
| Downregulated | **6** | **21.7** | **0** | **3.4** | 20 | 17.3 |

## Discussion

Here we show that male-limited X-chromosome evolution affected phenotypic traits and gene expression in a way that was mostly, but not entirely, consistent with our initial predictions. We expected to see an antagonistic change in sex-specific fitness, but this prediction was not borne out because although male fitness increased, we did not find any concomitant decrease in female fitness. However we did find evidence of an overall masculinization of locomotory activity, which is previously characterized as sexually antagonistic trait (Long and Rice 2007). We also found considerable evidence of a masculinization of the expression profile in the selected populations, and evidence that this response is in part a result of altered dosage compensation effects and release from mitonuclear conflict.

### Caveats

There are several caveats to these results that should be kept in mind, some of which serve to make our analysis more conservative. Our experimental protocol made it impossible to keep the effective population sizes of the X and autosomes equal between the control and selected treatments. We elected to reduce the effective population size of the X in the MLX treatment in order to avoid confounding differences in autosomal standing genetic variation with the selection treatment. This reduction in X chromosome population size should serve to limit the response to the selection treatment rather than enhance it. Similarly, due to logistical constraints at the time of expression data collection, we elected to use microarrays instead of RNAseq. RNAseq is superior for detecting low abundance transcripts, analysis of different isoforms, and generally produces lower amounts of technical variation compared to microarrays (Marioni et al. 2008, Daines et al. 2011). This implies that our results are more likely to suffer from low resolution rather than extensive false positives (unfortunately the lines are no longer available for complementary analysis using RNAseq for confirmation). Because changes in expression were investigated using whole-fly extractions rather than organs, we are also unable to distinguish between changes resulting from true upregulation versus changes in relative organ size (allometric effects). This limits our ability to determine exactly which mechanism(s) caused the changes in expression levels, but is sufficient for achieving our main aim of detecting evidence of changes associated with traits previously characterized as sexually antagonistic.

### Phenotypic data shows weak evidence of sexual antagonism

Several previous experiments have found that sex-limited selection leads to an increase in the fitness of the selected sex, and a decrease in the fitness of the unselected sex in this species (Rice 1992, Prasad et al. 2007, Morrow et al. 2008). Female-limited selection generally seems to have a smaller effect in the selected sex (~10% increase in Rice 1992 and Morrow et al. 2008) than male-limited selection (~15% increase in Prasad et al. 2007 and and Morrow et al. 2008). Interestingly, we found a larger increase in fitness in males (~25%) than in any of these previous studies. This is particularly surprising given that a smaller portion of the genome (i.e. the X) had the opportunity to respond to selection in this study compared to the whole-genome approaches of Prasad et al. (2007) and Morrow et al. (2008). This could indicate a large contribution of X-linked loci to male fitness, but may also be due to difference in number of generations (25-29 vs. >40). The usual explanation for an associated decrease in fitness in the unselected sex is sexual antagonism, although mutation accumulation at sex-limited loci is also a possibility in long-term experiments. We were therefore surprised that we did not recover any signal of sexual antagonism in fitness (Figure 2A).

One possibility is that antagonism had been resolved in this population at the time of data collection (Collet et al. 2016). However several lines of evidence suggest that antagonism may have existed but that we were unable to detect it here: other traits showed signatures of release from constraint imposed by selection in the other sex (e.g. locomotory activity and egg to adult survival, Figure 2B, Figures S2-3); there were signatures of phenotypic masculinization (Figure 2C); and changes in gene expression occurred in genes that were previously identified as sexually antagonistic (Table 3). There are then two plausible explanations for the somewhat puzzling lack of a decrease in female fitness. The first is that our experimental protocol for measuring female fitness did not capture all the relevant fitness variation. This is certainly possible, although fitness assays were carried out in such a way as to reflect fitness under the experimental culturing protocol and are similar to those used in previous studies of sexual antagonism in the population (Gibson et al. 2002, Prasad et al. 2007), and should therefore be relevant. However since any effects of the selection treatment on females were indirect, it is possible that there may have been mildly deleterious effects on female fitness that we did not have the power to detect, but which could have been better captured by investigating changes in fitness components that were not measured here (e.g. feeding efficiency or senescence).

The second possible explanation is dominance effects on the X-chromosome. It has been predicted that X-linked male-benefit/female-detriment alleles should preferentially be recessive in females, and female-benefit/male-detriment alleles should be dominant. Rice (1984) showed that both conditions allow high equilibrium frequencies of sexually antagonistic alleles. If most of the sexually antagonistic variation in the ancestral population was recessive in females, this could explain why there was no evidence of a reduction in female fitness (Figure 2A). The presence of a Control X chromosome in the MLX assay females would in this case mask both an increase in the frequency of recessive male-benefit alleles and a decrease in the frequency of dominant female-benefit alleles. These two explanations are not mutually exclusive, and both may have contributed to the lack of a decrease in female fitness. Although it is also possible that a signature of antagonism evolved between the time when the fitness data was collected (generation 40) and the RNA extractions were carried out (generation 50), we do not consider this explanation particularly likely since the other sex-limited evolution studies discussed above have detected signatures of antagonism after less than 30 generations (Rice 1992, Prasad et al. 2007, Morrow et al. 2008).

### Genomic location and function of genes that changed in expression

Genes that responded to the selection treatment were distributed throughout the genome (Table 1), with only around 80 of the transcripts that were identified as differentially expressed located on the X chromosome. This suggests that most of the changes that occurred are in genes that are regulated by X-linked loci, a handful of which might be sufficient to drive the observed downstream changes.

This is generally consistent with other results that suggest that sexually antagonistic loci may be few but of relatively large effect (Rice 1992, Barson et al. 2015, Ruzicka et al. 2019). Interestingly, downregulated transcripts were preferentially located on the X (Table 1). This is consistent with theory and previous empirical data that suggest that the X should be feminized (or at least demasculinized) compared to the autosomes (Rice 1984, Parisi et al. 2003, Sturgill et al. 2007, Long et al. 2012). The highly significant overrepresentation of upregulated transcripts on chromosome 4 (Table 1) may or may not be phenotypically relevant. Because chromosome 4 has very low rates of recombination, it is sufficient that a single gene which interacts with the X chromosome be selected for increased expression in order to cause a correlated response across many genes on chromosome 4. Nevertheless, chromosome 4 has been proposed to be the remnant of an old sex chromosome in *Drosophila* (Vicoso and Bachtrog 2013), so interactions between chromosome 4 and the X chromosome could therefore be functionally important. In addition, chromosome 4 has been shown to be disproportionately important in determining viability (Kenyon 1967, Charlesworth 2015), so changes in expression on chromosome 4 could be the mechanism behind the apparent increase in male survival seen here as a result of male-limited selection (Figure S2-S3). Finally, much of this overrepresentation may also be related to mitonuclear conflict, since 6 of the 16 upregulated genes located on chromosome 4 are previously characterized as mito-sensitive (i.e. influence male fitness depending on mitochondrial genotype).

The transcripts that changed in expression as a result of the selection treatment were in some cases consistent with previous phenotypic data, but were unexpected in other cases and suggest avenues for future exploration. A common theme within the overrepresented GO terms for upregulated genes was metabolism (Table S5). This is interesting because the direction of the change is consistent with increased adult activity levels (Figure 2B), which has previously been shown to be a sexually antagonistic trait (Long and Rice 2007), but is in the opposite direction to what we would expect from sexual dimorphism (males have a lower metabolic rate than females; Van Voorhies et al. 2004).

However this pattern could also be related to mitonuclear conflict, since mitochondrial genes preferentially accumulate male-deletrious alleles due to their female-limited transmission (Frank and Hurst 1996, Innocenti et al. 2011). The X chromosome has a reduced number of mito-sensitive genes compared to the autosomes in *Drosophila melanogaster*, suggesting the X is indeed a bad location for mitonuclear genes because selection against male-deleterious alleles can only occur via their indirect effect of reduced inheritance via the matriline (Rogell et al. 2014). However mito-sensitive genes are also overrepresented among genes important for male fitness (Rogell et al. 2014), so by decoupling inheritance of the X and the mitochondria, our selection treatment may have allowed more efficient selection against male-deleterious mito-associated alleles. Consistent with this, we found that both mito-annotated (Rogell et al. 2014) and mito-sensitive (Innocenti et al. 2011) genes were upregulated more often than expected in our dataset. There were also a number of terms associated with locomotory activity or muscle development, and testis-specific genes were overrepresented among upregulated genes, so another non-exclusive explanation could be that upregulation of metabolism is a result of increased overall activity levels or selection for improved performance in sperm competition.

Other interesting GO terms for upregulated genes include larval behaviour, many terms related to various aspects of vision (light stimulus, photoreceptor differentiation, phototaxis, and detection of visible light), and learning, memory and nervous system development (Table S5). Nervous tissue is energetically costly relative to other tissues, so this could also be a contributing factor to the observed effect on metabolic and mitonuclear genes discussed above. Since larval behaviour was not measured, it is unclear what changes may have occurred in this trait, but based on previous results a plausible explanation could be changes in feeding behaviour related to altered growth rates (Prasad et al. 2007). The large number of GO terms related to vision was unexpected, but could be a result of selection for increased mate searching efficiency or evaluation of competitors. Recent results suggest that vision may be more important to mating in *Drosophila* than was previously expected (Ribeiro et al. 2018). Similarly, learned male mate choice has been demonstrated in the ancestral population (Verzijden et al. 2015), and it is known that males use memory to evaluate their risk of sperm competition and allocate resources accordingly (Rouse et al. 2018), so terms related to learning, memory and nervous system development may also be a result of selection via mating interactions. The fact that there were a number of GO terms associated with male mating behaviour is consistent with mating interactions as the main mediator of the response to the selection treatment.

There were a number of genes specifically associated with oogenesis among downregulated genes (Table S6), which seems like a rather logical outcome. There were also very many GO terms associated with cell cycle regulation, DNA replication, and damage repair among downregulated genes. It is not clear what phenotypic traits these terms relate to, since they may be implicated in many different aspects of the phenotype. However we speculate that terms related to damage repair are a result of selection for decreased investment in somatic maintenance, since males have shorter longevity than females (Nuzhdin et al. 1997), and longevity has been shown to be a sexually antagonistic trait in several species, including *Drosophila* (Bonduriansky et al. 2008, Lewis et al. 2011, Berg and Maklakov 2012, Griffin et al. 2016). The overrepresentation of downregulated genes expressed in the crop would be consistent with this explanation, since the crop is a food storage organ. Terms related to various aspects of DNA transcription could be associated with altered dosage compensation effects (see below), while terms related to cell cycle regulation might be related to sperm production (Argyridou et al. 2017) or larval growth rate (Prasad et al. 2007).

### Sexual antagonism in gene expression and the evolution of sexual dimorphism

In contrast to results from phenotypic data, expression data showed a much stronger signature of sexual antagonism, and were consistent with expectations from extant sexual dimorphism. Genes that changed significantly in expression did so overwhelmingly in the same direction as extant sexual dimorphism (Figure 2), – i.e. genes that were already upregulated in males increased in expression, and genes that were downregulated decreased in expression – even though the magnitude of the change was small. In addition, genes previously identified as significantly male-biased in expression were overrepresented among upregulated transcripts, and female-biased genes were overrepresented among downregulated transcripts (Table 3), despite the fact that male-biased genes are generally underrepresented on the X chromosome (Parisi et al. 2003). These results are consistent with the prediction that sexual dimorphism is often a signature of fully or partially resolved sexual antagonism, even though the two need not always coincide (Cox and Calsbeek 2009, Innocenti and Morrow 2010). It also suggests that if X-linked expression could be fully decoupled between the sexes, then this would lead to an overall increase in the degree of sexual dimorphism.

Interestingly, there was a signature of antagonism among upregulated genes, since there was a strong relationship between previous characterization as being sexually antagonistic and upregulation after the selection treatment (Table 2). A caveat here is the fact that cryptic population substructure has since been found in the dataset used to determine the sexually antagonistic nature of these loci (M. Reuter, personal communication), and this may have inflated the signature of antagonism in Innocenti and Morrow (2010). Nevertheless, an overrepresentation of such loci among upregulated genes is consistent with our *a priori* predictions. There was no evidence of overrepresentation of loci more recently detected as sexually antagonistic (Ruzicka et al. 2019), but many of these loci were inferred to be coding changes unlikely to affect expression, and may therefore not be detectable within our dataset.

Increased sexual conflict has been suggested to induce an overall shift towards the male expression optimum (Hollis et al. 2014, Immonen et al. 2014, Innocenti et al. 2014, Perry et al. 2016), and might therefore be expected to parallel the changes seen here as a result of reducing female-specific selection and resolving sexual antagonism (but see Veltsos et al. 2017). Evidence for parallel changes was equivocal. There was no significant overrepresentation of genes identified by Innocenti el al. (2014), but there was overrepresentation of genes identified by Immonen et al. (2014). Given that changes as a result of alterations of sexual conflict seem to preferentially involve sex-biased genes (Hollis et al. 2014, Immonen et al. 2014, Innocenti et al. 2014), but that the direction of the change is sometimes opposite to the direction of sex-bias (Veltsos et al. 2017, Parker et al. 2019), it seems likely that any overlap with sexually antagonistic loci is caused by the fact that both phenomena are related to sexual dimorphism, and not because increased sexual conflict resolves antagonism *per se*.

Resolution of sexual antagonism on the X may partly be mediated by dosage compensation, since there was a positive relationship between degree of upregulation and distance from high-affinity sites (HAS), which are associated with dosage compensation (Figure 4). It is known from cross-species comparisons that that high-expression male-biased genes are more often located on the autosomes, while low-expression male biased genes are more often located on the X, and that most X-linked male-biased genes are located outside of dosage compensation regions (Bachtrog et al. 2010). The positive association seen here is consistent with previous results demonstrating that dosage compensation is a constraint for male-biased genes, although with the data at had we cannot determine the direction of causality – it could be that genes located farther from HAS changed more simply because they are more likely to be male-biased, not because genes located close to HAS were constrained in their response. However this result is also interesting because it suggests that there is standing genetic variation for the degree or consistency of dosage compensation, something that bears further investigation.

There has been some discussion and conflicting results reported as to whether the X should be feminized (enriched in female-biased genes compared to the autosomes; e.g. Parisi et al. 2003), demasculinized (impoverished for male-biased genes compared to the autosomes; e.g. Sturgill et al. 2007), both (reviewed in Dean and Mank 2014), or even masculinized (Patten 2018). Which pattern is most prevalent may depend not only on dominance but also the nature of sex-specific mutational effects (Frank and Patten 2019). Another complicating factor seems to be timescale. The X chromosome often accumulates mutations more quickly than the autosomes due to more efficient selection of beneficial mutations and/or drift, a phenomenon known as the faster-X effect. Since testis-specific genes have generally been shown to be rapidly evolving, a larger proportion of these rapidly accumulating mutations may be X-linked in origin. Indeed, Zhang *et al.* (2010) found that young male-biased genes were enriched on the X in *Drosophila*, but that old male-biased genes were enriched on the autosomes, consistent with the idea that the X contributes to rapid evolutionary change, but that it is an unfavourable location for male-biased genes. The microevolutionary effects seen in this study are consistent with these macroevolutionary patterns – when selection in females was removed, X-linked female-biased/female-benefit and male-detriment genes were downregulated. However by the same token, there were almost twice as many genes with significant upregulation compared to downregulation (342 vs. 176). This is consistent with the idea that X-linked male-biased genes are more constrained by selection in females, than female-biased genes are constrained by selection in males, and that these X-linked male-biased genes show a larger response when selection in females is removed.

### Conclusions

By forcing the X chromosome to only be expressed in males over many generations, we changed the selection pressures on the X to become similar to those experienced by the Y chromosome. Releasing males from constraints arising from counter-selection in females is predicted to lead to specialization for male fitness, and particularly to masculinization of phenotypes that normally experience sexually antagonistic selection. Indeed, we found evidence of masculinization primarily via upregulation of male-benefit genes, and downregulation of X-linked female benefit genes. In addition, we found evidence that female locomotory activity became masculinized, a trait that has previously been identified as sexually antagonistic. Changes in other traits not previously characterized as sexually antagonistic in this species, such as vision and learning/memory, suggest that these traits may be valuable to study further in this context in future. Interestingly, we could detect evidence of microevolutionary changes consistent with previously documented macroevolutionary patterns in sex chromosome evolution, such as upregulation of male-biased genes and downregulation of female-biased genes after a chromosome becomes male-limited, an increase in the expression of metabolic genes related to mitonuclear conflict, and evidence that dosage compensation effects can be rapidly altered. These results confirm the importance of the X in the evolution of sexual dimorphism and as a source for sexually antagonistic genetic variation, and demonstrate that experimental evolution can be a fruitful method for testing theories of sex chromosome evolution.

## Acknowledgements

This study was supported by the Swedish Research Council and Carl Tryggers Foundation (to JKA). Thanks to the SexGen group for comments on the manuscript. Thanks also to Björn Rogell for providing data for the analysis of mito-sensitive genes, and to Elina Immonen for providing data from Immonen et al. (2014).

