## Supplementary information for "The microevolutionary response to male-limited X-chromosome evolution in *Drosophila melanogaster* reflects macroevolutionary patterns"

### Supplementary methods

#### Fly stocks

All populations were derived from the $LH_M$ stock (Chippindale and Rice 2001), and were maintained so as to follow the ancestral culturing protocol. Sequencing data suggests that they are infected with *Wolbachia* (Gilks et al. 2016). $LH_M$ flies are fed with cornmeal-molasses-yeast medium and kept at 25°C, with a 12-12 light-dark cycle and 50% relative humidity. Adult females lay eggs on day zero, eggs hatch on day 1, and adult flies eclose on day 9 or 10. After eclosion, males and females are mixed between vials on day 12, and 16 randomly-selected pairs are placed in each vial to produce the next generation. Males and females are then allowed 48 hours to interact (primarily via mating and competition for yeast) in vials supplemented with approximately 6 mg of live yeast, after which they are transferred to new unyeasted vials for oviposition. Flies are kept in mixed-sex groups in the new vials, and females have an 18 hour window in which they can lay eggs. All adult flies are discarded at the end of this period (new day 1). Vials are trimmed to 150 ± 10 eggs in order to keep larval densities constant. A total of 56 $LH_M$ stock vials are maintained (1 792 breeding adults and about 8 000 juveniles), preserving a large population size with substantial standing genetic variation (Gilks et al. 2016). This culturing protocol ensures that flies are given ample opportunity to interact with unrelated individuals from other vials, as well as strictly controlling their density at both adult and larval stages.

#### Recombination box

The role of the recombination box was to allow low levels of recombination so that the MLX selection treatment would not be compromised by hitchhiking of deleterious alleles. Each recombination box was kept at 10% of the MLX population size, which should be enough to override any effects of genetic hitchhiking of deleterious alleles (Rice 1998). In each generation new males for the recombination box were randomly selected from the paired MLX experimental population, and combined with females from the recombination box. An equal number of recombination box males were returned to the main MLX population. This results in a constant inflow of new X-chromosomes to the recombination box, and an outflow of recombined X-chromosomes from the recombination box back into the main MLX populations (for details, see [citation suppressed to maintain anonymity]).

#### Standard fitness assays

Wildtype red-eyed flies from the experimental populations are the target individuals, and the competitors are an $LH_M$-derived population that has had a recessive brown eye-colour marker (*$bw^1$*) introgressed. The standard culturing protocol described above is followed, but with two exceptions: 1. On day 12, 4 target flies of the desired sex are combined with 12 brown-eyed individuals of the same sex and 16 brown-eyed individuals of the opposite sex. 2. Instead of transferring all flies to new vials for oviposition 48 hours later, females are moved to individual test tubes for egg laying. Males are discarded when the females are moved, and females are discarded after 18 hours of oviposition. Female fitness is measured as the number of eggs laid per test tube, and is averaged within a vial since females from the same vial are not independent of each other. Male fitness is measured as the proportion of adult offspring sired compared to the competitor males. Again, target males from the same vial are not independent, but for the male data results were pooled within a vial, since the progeny of individual target males cannot be identified.

#### Network analyses

Two interaction network analyses were also carried out, firstly to investigate degree of overlap with a previous dataset, and secondly to determine whether most significant genes fall into several large or many small clusters. The first analysis was of first-order interactions within the genes identified as having significantly altered expression in this study using the esyN online network tool (Bean et al. 2014). This analysis was to determine whether most genes that changed participated in the same networks or not. Parts of this dataset have already been analysed elsewhere (Abbott et al. 2013), so we also checked for overlap among genes identified as significantly different in expression in both datasets, and whether the estimated change was similar or not. Very few genes were determined to be significant in both datasets, so we generated an interaction network for genes in both sets using the online tool NetVenn (Wang et al. 2014). We did this in order to determine whether the lack of overlap was because the genes were part of the same interaction networks, but that which ones come up as stastistically significant is to some dependent on chance, as has previously been shown in *Drosophila melanogaster* (Huang et al. 2012).

### Supplementary results

#### Phenotypic assays

Table S1: Results from a factorial anova of the effect of sex and selection treatment on relative fitness. There is a significant effect of selection treatment only.

| Effect | DF | MS | *F* | *P*-value |
|---|---|---|---|---|
| **Sex** | 1 | 0 | 0.0000 | 1.0000 |
| **Selection** | 1 | 0.0797 | 6.1088 | ***0.0386*** |
| **Sex*Selection** | 1 | 0.0147 | 1.1247 | 0.3199 |
| **Error** | 8 | 0.0130 | | |

Table S2: Results of generalized linear models of the effect of (a) sex and selection treatment on sex ratio, (b) selection treatment on egg to adult survival, and (c) sex and selection treatment on locomotory activity. All effects are significant, except for the main effect of selection treatment on locomotory activity.

| Phenotype | Effect | DF | Deviance | Residual DF | Residual deviance | *P*-value |
|---|---|---|---|---|---|---|
| (a) **Sex Ratio** | Residual | | | 11 | 0.40735 | |
| | Sex | 1 | 0.137065 | 10 | 0.27029 | ***$2.686*10^{-8}$*** |
| | Selection | 1 | 0.185573 | 9 | 0.08472 | ***$9.775*10^{-11}$*** |
| | Sex*Selection | 1 | 0.049292 | 8 | 0.03542 | ***0.0008538*** |
| (b) **Survival** | Residual | | | 5 | 0.83699 | |
| | Selection | 1 | 0.6748 | 4 | 0.16218 | ***$5.409*10^{-05}$*** |
| **(c) Locomotory activity** | Residual | | | 11 | 0.97778 | |
| | Sex | 1 | 0.44870 | 10 | 0.52909 | ***$1.686*10^{-06}$*** |
| | Selection | 1 | 0.06338 | 9 | 0.46571 | 0.07195 |
| | Sex*Selection | 1 | 0.30828 | 8 | 0.15743 | ***$7.230*10^{-05}$*** |

Table S3: Results from nested random effect models of the effect of sex and selection treatment on (a) relative fitness, (b) sex ratio, (c) survival, and (d) locomotory activity. Fitness measures were converted to Z-scores before analysis in order to avoid violating assumptions of the *F*-tests. Note that it was not necessary to carry out generalized linear mixed models for sex ratio, survival and locomotory activity since the nested random effect models adequately fulfilled the assumptions of the *F*-tests. For sex ratio, *F*-tests were obtained using type 3 sums of squares and Satterthwaite's approximation for denominator degrees of freedom. All results are qualitatively similar to the means-based analyses except for fitness, and the lack of a significant interaction effect on sex ratio (compare with Tables S1 and S2).

| **Phenotype** | **Effect** | **NumDF** | **DenDF** | **MS** | ***F*** | ***P*-value** | **Variance** |
|---|---|---|---|---|---|---|---|
| (a**) Fitness** | Sex | 1 | 112 | 0 | 0.0000 | 1.0000 | - |
| | Selection | 1 | 4 | 1.04044 | 1.1261 | 0.3484 | - |
| | Sex*Selection | 1 | 112 | 0.01483 | 0.0161 | 0.8994 | - |
| | Population(Group) | - | - | - | - | - | 0.09529 |
| | Residual | - | - | - | - | - | 0.92393 |
| **(b) Sex ratio** | Sex | 1 | 110 | 0.29164 | 8.8340 | ***0.0036*** | - |
| | Selection | 1 | 14.96 | 0.38973 | 10.789 | ***0.0050*** | - |
| | Sex*Selection | 1 | 110.07 | 0.06828 | 1.8452 | 0.1771 | - |
| | Population(Group) | - | - | - | - | - | 0.000 |
| | Residual | - | - | - | - | - | 0.037 |
| **(c) Survival** | Selection | 1 | 4 | 0.51063 | 18.173 | ***0.0130*** | - |
| | Population(Group) | - | - | - | - | - | 0.00630 |
| | Residual | - | - | - | - | - | 0.02810 |
| **(d) Locomotory activity** | Sex | 1 | 4 | 0.48600 | 0.0000 | ***0.0001*** | - |
| | Selection | 1 | 52 | 0.05835 | 1.1261 | 0.2204 | - |
| | Sex*Selection | 1 | 52 | 0.26667 | 0.0161 | ***0.0031*** | - |
| | Population(Group) | - | - | - | - | - | 0.00039 |
| | Residual | - | - | - | - | - | 0.02772 |

#### Up- versus downregulated transcripts

Table S4: Tissue-specificity of MLX transcripts. Upregulated transcripts are previously characterized as being differentially expressed in several tissues more often than expected. Downregulated transcripts are only expressed in the crop more often than expected by chance. Significant values are highlighted.

| Tissue | Up-regulated transcripts | | Down-regulated transcripts | |
|---|---|---|---|---|
| | Odds ratio | Adjusted *P*-value | Odds ratio | Adjusted *P*-value |
| **Testes** | ***2.5034*** | ***0.004711*** | 0.2187 | 1 |
| **Accessory gland** | 0.9803 | 1 | 0.1799 | 1 |
| **Ejaculatory duct** | 1.2395 | 1 | 0.2156 | 1 |
| **Ovary** | ***2.9917*** | $\boldsymbol{5.27*10^{-11}}$ | 0.2254 | 1 |
| **Virgin spermatheca** | ***6.5911*** | $\boldsymbol{2.02*10^{-47}}$ | 0.1912 | 1 |
| **Mated spermatheca** | ***9.3661*** | $\boldsymbol{6.14*10^{-82}}$ | 0.2090 | 1 |
| **Crop** | 0.1823 | 1 | ***18.1912*** | $\boldsymbol{2.91*10^{-70}}$ |
| **Mid gut** | 1.4764 | 0.520739 | 0.2569 | 1 |
| **Hind gut** | ***10.3167*** | $\boldsymbol{2.84*10^{-86}}$ | 0.2531 | 1 |
| **Thoracic ganglion** | 0 | 1 | 0 | 1 |
| **Fat body** | ***5.0144*** | $\boldsymbol{5.02*10^{-29}}$ | 0.3812 | 1 |
| **Heart** | ***3.0259*** | $\boldsymbol{2.36*10^{-12}}$ | 0.5404 | 1 |
| **Head** | 0.6891 | 1 | 0.0634 | 1 |
| **Brain** | 1.4161 | 1 | 0.1177 | 1 |

Table S5: Overrepresented Gene Ontology database terms for transcripts that were up-regulated in MLX individuals relative to controls. Transcripts are associated with, among other things, synaptic transmission, muscle development, metabolism, and locomotory behaviour.

| Domain | ID | P-value | Odds ratio | Expected count | Observed count | Size | Term |
|---|---|---|---|---|---|---|---|
| Biological process | GO:0007268 | 0 | 12.829 | 5 | 31 | 71 | Synaptic transmission |
| Biological process | GO:0019226 | 0 | 12.513 | 5 | 31 | 72 | Transmission of nerve impulse |
| Biological process | GO:0003008 | 0 | 5.99 | 12 | 48 | 188 | System process |
| Biological process | GO:0007267 | 0 | 9.817 | 6 | 32 | 86 | Cell-cell signaling |
| Biological process | GO:0001505 | 0 | 18.506 | 3 | 22 | 41 | Regulation of neurotransmitter levels |
| Biological process | GO:0032501 | 0 | 2.852 | 65 | 119 | 1009 | Multicellular organismal process |
| Biological process | GO:0050877 | 0 | 5.253 | 11 | 41 | 173 | Neurological system process |
| Biological process | GO:0065008 | 0 | 4.096 | 17 | 50 | 261 | Regulation of biological quality |
| Biological process | GO:0007154 | 0 | 4.434 | 14 | 45 | 218 | Cell communication |
| Biological process | GO:0007269 | 0 | 16.635 | 2 | 18 | 35 | Neurotransmitter secretion |
| Biological process | GO:0023052 | 0 | 3.089 | 34 | 76 | 531 | Signaling |
| Biological process | GO:0003001 | 0 | 14.875 | 2 | 18 | 37 | Generation of a signal involved in cell-cell signaling |
| Biological process | GO:0023046 | 0 | 3.251 | 26 | 62 | 400 | Signaling process |
| Biological process | GO:0023060 | 0 | 3.251 | 26 | 62 | 400 | Signal transmission |
| Biological process | GO:0032940 | 0 | 12.836 | 3 | 18 | 40 | Secretion by cell |
| Biological process | GO:0046903 | 0 | 11.502 | 3 | 19 | 45 | Secretion |
| Biological process | GO:0007610 | 0 | 4.448 | 12 | 38 | 180 | Behavior |
| Biological process | GO:0006836 | 0 | 9.33 | 3 | 19 | 51 | Neurotransmitter transport |
| Biological process | GO:0048489 | 0 | 16.646 | 2 | 14 | 27 | Synaptic vesicle transport |
| Biological process | GO:0055001 | 0 | 13.513 | 2 | 14 | 30 | Muscle cell development |
| Biological process | GO:0055002 | 0 | 13.513 | 2 | 14 | 30 | Striated muscle cell development |

| | | | | | | | |
|---|---|---|---|---|---|---|---|
| Biological process | GO:0065007 | 0 | 2.248 | 71 | 114 | 1113 | Biological regulation |
| Biological process | GO:0051146 | 0 | 9.673 | 2 | 15 | 39 | Striated muscle cell differentiation |
| Biological process | GO:0048731 | 0 | 2.461 | 38 | 72 | 590 | System development |
| Biological process | GO:0007626 | 0 | 6.036 | 5 | 20 | 72 | Locomotory behavior |
| Biological process | GO:0006887 | 0 | 16.8 | 1 | 11 | 21 | Exocytosis |
| Biological process | GO:0006811 | 0 | 4.085 | 9 | 29 | 143 | Ion transport |
| Biological process | GO:0042692 | 0 | 8.591 | 3 | 15 | 42 | Muscle cell differentiation |
| Biological process | GO:0016079 | 0 | 19.02 | 1 | 10 | 18 | Synaptic vesicle exocytosis |
| Biological process | GO:0009888 | 0 | 3.447 | 12 | 34 | 194 | Tissue development |
| Biological process | GO:0006812 | 0 | 4.596 | 7 | 23 | 102 | Cation transport |
| Biological process | GO:0006810 | 0 | 2.286 | 41 | 74 | 644 | Transport |
| Biological process | GO:0006873 | 0 | 15.208 | 1 | 10 | 20 | Cellular ion homeostasis |
| Biological process | GO:0051234 | 0 | 2.197 | 43 | 75 | 674 | Establishment of localization |
| Biological process | GO:0007275 | 0 | 2.134 | 48 | 81 | 754 | Multicellular organismal development |
| Biological process | GO:0014706 | 0 | 10.482 | 2 | 11 | 27 | Striated muscle tissue development |
| Biological process | GO:0060537 | 0 | 10.482 | 2 | 11 | 27 | Muscle tissue development |
| Biological process | GO:0007519 | 0 | 12.666 | 1 | 10 | 22 | Skeletal muscle tissue development |
| Biological process | GO:0050801 | 0 | 12.666 | 1 | 10 | 22 | Ion homeostasis |
| Biological process | GO:0055082 | 0 | 12.666 | 1 | 10 | 22 | Cellular chemical homeostasis |
| Biological process | GO:0048856 | 0 | 2.122 | 46 | 78 | 724 | Anatomical structure development |
| Biological process | GO:0006874 | 0 | 35.107 | 1 | 7 | 10 | Cellular calcium ion homeostasis |
| Biological process | GO:0006875 | 0 | 35.107 | 1 | 7 | 10 | Cellular metal ion homeostasis |
| Biological process | GO:0055074 | 0 | 35.107 | 1 | 7 | 10 | Calcium ion homeostasis |
| Biological process | GO:0030001 | 0 | 5.378 | 4 | 17 | 66 | Metal ion transport |

| | | | | | | | |
|---|---|---|---|---|---|---|---|
| Biological process | GO:0051179 | 0 | 2.076 | 51 | 83 | 792 | Localization |
| Biological process | GO:0055065 | 0 | 26.323 | 1 | 7 | 11 | Metal ion homeostasis |
| Biological process | GO:0060538 | 0 | 8.82 | 2 | 11 | 30 | Skeletal muscle organ development |
| Biological process | GO:0048513 | 0 | 2.261 | 30 | 56 | 471 | Organ development |
| Biological process | GO:0008344 | 0 | 11.351 | 1 | 9 | 21 | Adult locomotory behavior |
| Biological process | GO:0030534 | 0 | 6.784 | 2 | 12 | 39 | Adult behavior |
| Biological process | GO:0048878 | 0 | 8.928 | 2 | 10 | 27 | Chemical homeostasis |
| Biological process | GO:0061061 | 0 | 4.043 | 6 | 19 | 92 | Muscle structure development |
| Biological process | GO:0048747 | 0 | 13.408 | 1 | 8 | 17 | Muscle fiber development |
| Biological process | GO:0032502 | 0 | 1.914 | 56 | 86 | 875 | Developmental process |
| Biological process | GO:0006816 | 0 | 12.064 | 1 | 8 | 18 | Calcium ion transport |
| Biological process | GO:0051239 | 0 | 3.068 | 10 | 25 | 153 | Regulation of multicellular organismal process |
| Biological process | GO:0019725 | 0 | 5.902 | 3 | 12 | 43 | Cellular homeostasis |
| Biological process | GO:0030003 | 0 | 15.029 | 1 | 7 | 14 | Cellular cation homeostasis |
| Biological process | GO:0030005 | 0 | 15.029 | 1 | 7 | 14 | Cellular di-, tri-valent inorganic cation homeostasis |
| Biological process | GO:0009605 | 0 | 3.867 | 6 | 18 | 90 | Response to external stimulus |
| Biological process | GO:0070838 | 0 | 10.964 | 1 | 8 | 19 | Divalent metal ion transport |
| Biological process | GO:0007399 | 0 | 2.335 | 21 | 41 | 325 | Nervous system development |
| Biological process | GO:0009653 | 0 | 2.002 | 35 | 59 | 549 | Anatomical structure morphogenesis |
| Biological process | GO:0007498 | 0 | 5.377 | 3 | 12 | 46 | Mesoderm development |
| Biological process | GO:0055066 | 0 | 11.683 | 1 | 7 | 16 | Di-, tri-valent inorganic cation homeostasis |
| Biological process | GO:0055080 | 0 | 11.683 | 1 | 7 | 16 | Cation homeostasis |

| | | | | | | | |
|---|---|---|---|---|---|---|---|
| Biological process | GO:0048869 | 0 | 1.982 | 33 | 56 | 522 | Cellular developmental process |
| Biological process | GO:0007186 | 0 | 3.637 | 6 | 17 | 89 | G-protein coupled receptor protein signaling pathway |
| Biological process | GO:0016192 | 0 | 2.939 | 9 | 23 | 145 | Vesicle-mediated transport |
| Biological process | GO:0015674 | 0 | 8.607 | 1 | 8 | 22 | Di-, tri-valent inorganic cation transport |
| Biological process | GO:0030154 | 0 | 2.018 | 30 | 52 | 474 | Cell differentiation |
| Biological process | GO:0050793 | 0 | 2.817 | 10 | 24 | 157 | Regulation of developmental process |
| Biological process | GO:0007517 | 0 | 3.896 | 5 | 15 | 74 | Muscle organ development |
| Biological process | GO:0003012 | 0 | 24.869 | 1 | 5 | 8 | Muscle system process |
| Biological process | GO:0006936 | 0 | 24.869 | 1 | 5 | 8 | Muscle contraction |
| Biological process | GO:0006813 | 0 | 8.031 | 1 | 8 | 23 | Potassium ion transport |
| Biological process | GO:0032989 | 0 | 2.271 | 18 | 35 | 280 | Cellular component morphogenesis |
| Biological process | GO:0030239 | 0 | 12.829 | 1 | 6 | 13 | Myofibril assembly |
| Biological process | GO:0042133 | 0 | 59.471 | 0 | 4 | 5 | Neurotransmitter metabolic process |
| Biological process | GO:0015672 | 0 | 3.75 | 5 | 14 | 71 | Monovalent inorganic cation transport |
| Biological process | GO:0007528 | 0 | 11.222 | 1 | 6 | 14 | Neuromuscular junction development |
| Biological process | GO:0030537 | 0 | 11.222 | 1 | 6 | 14 | Larval behavior |
| Biological process | GO:0048741 | 0 | 11.222 | 1 | 6 | 14 | Skeletal muscle fiber development |
| Biological process | GO:0050896 | 0 | 1.905 | 32 | 52 | 496 | Response to stimulus |
| Biological process | GO:0050808 | 0 | 6.687 | 2 | 8 | 26 | Synapse organization |
| Biological process | GO:0048468 | 0 | 2.004 | 25 | 43 | 387 | Cell development |
| Biological process | GO:0050803 | 0 | 9.972 | 1 | 6 | 15 | Regulation of synapse structure and activity |

| | | | | | | | |
|---|---|---|---|---|---|---|---|
| Biological process | GO:0045214 | 0 | 29.727 | 0 | 4 | 6 | Sarcomere organization |
| Biological process | GO:0007507 | 0 | 6.015 | 2 | 8 | 28 | Heart development |
| Biological process | GO:0008291 | 0 | Inf | 0 | 3 | 3 | Acetylcholine metabolic process |
| Biological process | GO:0042136 | 0 | Inf | 0 | 3 | 3 | Neurotransmitter biosynthetic process |
| Biological process | GO:0090130 | 0 | 8.973 | 1 | 6 | 16 | Tissue migration |
| Biological process | GO:0051649 | 0 | 2.326 | 13 | 26 | 200 | Establishment of localization in cell |
| Biological process | GO:0055085 | 0 | 2.312 | 13 | 26 | 201 | Transmembrane transport |
| Biological process | GO:0048699 | 0 | 2.201 | 15 | 29 | 235 | Generation of neurons |
| Biological process | GO:0042592 | 0 | 3.467 | 4 | 13 | 70 | Homeostatic process |
| Biological process | GO:0050789 | 0 | 1.62 | 65 | 89 | 1022 | Regulation of biological process |
| Biological process | GO:0007218 | 0 | 6.559 | 1 | 7 | 23 | Neuropeptide signaling pathway |
| Biological process | GO:0031032 | 0 | 6.559 | 1 | 7 | 23 | Actomyosin structure organization |
| Biological process | GO:0042330 | 0 | 6.559 | 1 | 7 | 23 | Taxis |
| Biological process | GO:0043062 | 0 | 4.315 | 3 | 10 | 45 | Extracellular structure organization |
| Biological process | GO:0007417 | 0 | 3.093 | 6 | 15 | 89 | Central nervous system development |
| Biological process | GO:0042221 | 0 | 2.511 | 10 | 21 | 150 | Response to chemical stimulus |
| Biological process | GO:0030182 | 0 | 2.224 | 14 | 27 | 216 | Neuron differentiation |
| Biological process | GO:0016545 | 0 | 19.813 | 0 | 4 | 7 | Male courtship behavior, veined wing vibration |
| Biological process | GO:0045433 | 0 | 19.813 | 0 | 4 | 7 | Male courtship behavior, veined wing generated song production |
| Biological process | GO:0048065 | 0 | 19.813 | 0 | 4 | 7 | Male courtship behavior, veined wing extension |
| Biological process | GO:0050794 | 0.001 | 1.602 | 60 | 82 | 938 | Regulation of cellular process |

| | | | | | | | |
|---|---|---|---|---|---|---|---|
| Biological process | GO:0022008 | 0.001 | 2.083 | 16 | 29 | 246 | Neurogenesis |
| Biological process | GO:0000902 | 0.001 | 2.105 | 15 | 28 | 235 | Cell morphogenesis |
| Biological process | GO:0007427 | 0.001 | 9.313 | 1 | 5 | 13 | Epithelial cell migration, open tracheal system |
| Biological process | GO:0010631 | 0.001 | 9.313 | 1 | 5 | 13 | Epithelial cell migration |
| Biological process | GO:0090132 | 0.001 | 9.313 | 1 | 5 | 13 | Epithelium migration |
| Biological process | GO:0007619 | 0.001 | 5.519 | 2 | 7 | 26 | Courtship behavior |
| Biological process | GO:0008345 | 0.001 | 14.855 | 1 | 4 | 8 | Larval locomotory behavior |
| Biological process | GO:0007270 | 0.001 | 44.42 | 0 | 3 | 4 | Nerve-nerve synaptic transmission |
| Biological process | GO:0035072 | 0.001 | 44.42 | 0 | 3 | 4 | Ecdysone-mediated induction of salivary gland cell autophagic cell death |
| Biological process | GO:0042044 | 0.001 | 44.42 | 0 | 3 | 4 | Fluid transport |
| Biological process | GO:0042439 | 0.001 | 44.42 | 0 | 3 | 4 | Ethanolamine and derivative metabolic process |
| Biological process | GO:0051208 | 0.001 | 44.42 | 0 | 3 | 4 | Sequestering of calcium ion |
| Biological process | GO:0051238 | 0.001 | 44.42 | 0 | 3 | 4 | Sequestering of metal ion |
| Biological process | GO:0009416 | 0.001 | 3.677 | 3 | 10 | 51 | Response to light stimulus |
| Biological process | GO:0007165 | 0.001 | 1.886 | 21 | 35 | 326 | Signal transduction |
| Biological process | GO:0007242 | 0.001 | 2.212 | 11 | 22 | 175 | Intracellular signaling cascade |
| Biological process | GO:0044057 | 0.002 | 5.973 | 1 | 6 | 21 | Regulation of system process |
| Biological process | GO:0009453 | 0.002 | 11.881 | 1 | 4 | 9 | Energy taxis |
| Biological process | GO:0042331 | 0.002 | 11.881 | 1 | 4 | 9 | Phototaxis |
| Biological process | GO:0031175 | 0.002 | 2.346 | 9 | 19 | 143 | Neuron projection development |
| Biological process | GO:0048812 | 0.002 | 2.346 | 9 | 19 | 143 | Neuron projection morphogenesis |
| Biological process | GO:0001667 | 0.002 | 7.446 | 1 | 5 | 15 | Ameboidal cell migration |

| | | | | | | | |
|---|---|---|---|---|---|---|---|
| Biological process | GO:0007409 | 0.002 | 2.724 | 6 | 14 | 92 | Axonogenesis |
| Biological process | GO:0030030 | 0.002 | 2.168 | 11 | 22 | 178 | Cell projection organization |
| Biological process | GO:0009887 | 0.002 | 1.94 | 17 | 29 | 261 | Organ morphogenesis |
| Biological process | GO:0048858 | 0.002 | 2.219 | 10 | 20 | 158 | Cell projection morphogenesis |
| Biological process | GO:0051641 | 0.002 | 1.97 | 15 | 27 | 239 | Cellular localization |
| Biological process | GO:0007274 | 0.002 | 22.204 | 0 | 3 | 5 | Neuromuscular synaptic transmission |
| Biological process | GO:0035078 | 0.002 | 22.204 | 0 | 3 | 5 | Induction of programmed cell death by ecdysone |
| Biological process | GO:0007411 | 0.002 | 3.073 | 4 | 11 | 65 | Axon guidance |
| Biological process | GO:0007166 | 0.003 | 1.95 | 15 | 27 | 241 | Cell surface receptor linked signaling pathway |
| Biological process | GO:0002165 | 0.003 | 2.146 | 11 | 21 | 171 | Instar larval or pupal development |
| Biological process | GO:0032990 | 0.003 | 2.17 | 10 | 20 | 161 | Cell part morphogenesis |
| Biological process | GO:0046530 | 0.003 | 2.833 | 5 | 12 | 76 | Photoreceptor cell differentiation |
| Biological process | GO:0023033 | 0.003 | 1.931 | 16 | 27 | 243 | Signaling pathway |
| Biological process | GO:0009314 | 0.003 | 3.135 | 4 | 10 | 58 | Response to radiation |
| Biological process | GO:0008049 | 0.003 | 6.202 | 1 | 5 | 17 | Male courtship behavior |
| Biological process | GO:0009584 | 0.003 | 6.202 | 1 | 5 | 17 | Detection of visible light |
| Biological process | GO:0060179 | 0.003 | 6.202 | 1 | 5 | 17 | Male mating behavior |
| Biological process | GO:0006928 | 0.003 | 2.231 | 9 | 18 | 141 | Cellular component movement |
| Biological process | GO:0007617 | 0.003 | 4.187 | 2 | 7 | 32 | Mating behavior |
| Biological process | GO:0001751 | 0.003 | 2.909 | 4 | 11 | 68 | Compound eye photoreceptor cell differentiation |
| Biological process | GO:0009791 | 0.004 | 2.074 | 11 | 21 | 176 | Post-embryonic development |
| Biological process | GO:0048667 | 0.004 | 2.212 | 9 | 18 | 142 | Cell morphogenesis involved in neuron differentiation |

| | | | | | | | |
|---|---|---|---|---|---|---|---|
| Biological process | GO:0001754 | 0.004 | 2.858 | 4 | 11 | 69 | Eye photoreceptor cell differentiation |
| Biological process | GO:0007522 | 0.004 | Inf | 0 | 2 | 2 | Visceral muscle development |
| Biological process | GO:0007637 | 0.004 | Inf | 0 | 2 | 2 | Proboscis extension reflex |
| Biological process | GO:0008292 | 0.004 | Inf | 0 | 2 | 2 | Acetylcholine biosynthetic process |
| Biological process | GO:0017157 | 0.004 | Inf | 0 | 2 | 2 | Regulation of exocytosis |
| Biological process | GO:0051780 | 0.004 | Inf | 0 | 2 | 2 | Behavioral response to nutrient |
| Biological process | GO:0060004 | 0.004 | Inf | 0 | 2 | 2 | Reflex |
| Biological process | GO:0007611 | 0.004 | 4.025 | 2 | 7 | 33 | Learning or memory |
| Biological process | GO:0006164 | 0.004 | 3.008 | 4 | 10 | 60 | Purine nucleotide biosynthetic process |
| Biological process | GO:0040011 | 0.004 | 2.369 | 7 | 15 | 111 | Locomotion |
| Biological process | GO:0019932 | 0.004 | 5.723 | 1 | 5 | 18 | Second-messenger-mediated signaling |
| Biological process | GO:0007629 | 0.004 | 14.798 | 0 | 3 | 6 | Flight behavior |
| Biological process | GO:0035081 | 0.004 | 14.798 | 0 | 3 | 6 | Induction of programmed cell death by hormones |
| Biological process | GO:0042391 | 0.004 | 14.798 | 0 | 3 | 6 | Regulation of membrane potential |
| Biological process | GO:0007635 | 0.005 | 3.423 | 3 | 8 | 43 | Chemosensory behavior |
| Biological process | GO:0006163 | 0.005 | 2.89 | 4 | 10 | 62 | Purine nucleotide metabolic process |
| Biological process | GO:0035075 | 0.005 | 7.419 | 1 | 4 | 12 | Response to ecdysone |
| Biological process | GO:0042398 | 0.005 | 7.419 | 1 | 4 | 12 | Cellular amino acid derivative biosynthetic process |
| Biological process | GO:0048545 | 0.005 | 7.419 | 1 | 4 | 12 | Response to steroid hormone stimulus |
| Biological process | GO:0009165 | 0.005 | 2.715 | 5 | 11 | 72 | Nucleotide biosynthetic process |
| Biological process | GO:0007416 | 0.006 | 5.313 | 1 | 5 | 19 | Synapse assembly |
| Biological process | GO:0008355 | 0.006 | 5.313 | 1 | 5 | 19 | Olfactory learning |
| Biological process | GO:0030036 | 0.006 | 2.67 | 5 | 11 | 73 | Actin cytoskeleton organization |

| | | | | | | | |
|---|---|---|---|---|---|---|---|
| Biological process | GO:0000904 | 0.007 | 2.074 | 10 | 18 | 150 | Cell morphogenesis involved in differentiation |
| Biological process | GO:0030029 | 0.007 | 2.627 | 5 | 11 | 74 | Actin filament-based process |
| Biological process | GO:0009628 | 0.007 | 2.393 | 6 | 13 | 95 | Response to abiotic stimulus |
| Biological process | GO:0048666 | 0.007 | 1.966 | 11 | 20 | 175 | Neuron development |
| Biological process | GO:0032879 | 0.007 | 2.933 | 4 | 9 | 55 | Regulation of localization |
| Biological process | GO:0008045 | 0.007 | 6.593 | 1 | 4 | 13 | Motor axon guidance |
| Biological process | GO:0016202 | 0.007 | 6.593 | 1 | 4 | 13 | Regulation of striated muscle tissue development |
| Biological process | GO:0050908 | 0.007 | 6.593 | 1 | 4 | 13 | Detection of light stimulus involved in visual perception |
| Biological process | GO:0045595 | 0.007 | 2.364 | 6 | 13 | 96 | Regulation of cell differentiation |
| Biological process | GO:0007584 | 0.007 | 11.096 | 0 | 3 | 7 | Response to nutrient |
| Biological process | GO:0048589 | 0.008 | 3.15 | 3 | 8 | 46 | Developmental growth |
| Biological process | GO:0051705 | 0.008 | 3.15 | 3 | 8 | 46 | Behavioral interaction between organisms |
| Biological process | GO:0006754 | 0.008 | 3.484 | 2 | 7 | 37 | ATP biosynthetic process |
| Biological process | GO:0045165 | 0.008 | 2.182 | 8 | 15 | 119 | Cell fate commitment |
| Biological process | GO:0007155 | 0.008 | 2.545 | 5 | 11 | 76 | Cell adhesion |
| Biological process | GO:0022610 | 0.008 | 2.545 | 5 | 11 | 76 | Biological adhesion |
| Biological process | GO:0010646 | 0.009 | 2.01 | 10 | 18 | 154 | Regulation of cell communication |
| Biological process | GO:0009583 | 0.009 | 3.887 | 2 | 6 | 29 | Detection of light stimulus |
| Biological process | GO:0019098 | 0.009 | 3.371 | 2 | 7 | 38 | Reproductive behavior |
| Biological process | GO:0043085 | 0.009 | 3.371 | 2 | 7 | 38 | Positive regulation of catalytic activity |
| Biological process | GO:0046034 | 0.009 | 3.371 | 2 | 7 | 38 | ATP metabolic process |
| Biological process | GO:0035023 | 0.01 | 5.932 | 1 | 4 | 14 | Regulation of Rho protein signal transduction |

| | | | | | | | |
|---|---|---|---|---|---|---|---|
| Biological process | GO:0048634 | 0.01 | 5.932 | 1 | 4 | 14 | Regulation of muscle organ development |
| Biological process | GO:0050962 | 0.01 | 5.932 | 1 | 4 | 14 | Detection of light stimulus involved in sensory perception |
| Biological process | GO:0006793 | 0.01 | 1.733 | 17 | 27 | 266 | Phosphorus metabolic process |
| Biological process | GO:0006796 | 0.01 | 1.733 | 17 | 27 | 266 | Phosphate metabolic process |
| Biological process | GO:0040007 | 0.01 | 2.468 | 5 | 11 | 78 | Growth |
| Cellular component | GO:0045202 | 0 | 11.185 | 4 | 23 | 56 | Synapse |
| Cellular component | GO:0043292 | 0 | 49.987 | 1 | 13 | 17 | Contractile fiber |
| Cellular component | GO:0044449 | 0 | 61.232 | 1 | 12 | 15 | Contractile fiber part |
| Cellular component | GO:0044456 | 0 | 11.973 | 3 | 19 | 44 | Synapse part |
| Cellular component | GO:0030016 | 0 | 50.517 | 1 | 10 | 13 | Myofibril |
| Cellular component | GO:0030017 | 0 | 50.517 | 1 | 10 | 13 | Sarcomere |
| Cellular component | GO:0016020 | 0 | 2.396 | 53 | 91 | 825 | Membrane |
| Cellular component | GO:0008021 | 0 | 16.716 | 1 | 11 | 21 | Synaptic vesicle |
| Cellular component | GO:0030136 | 0 | 16.716 | 1 | 11 | 21 | Clathrin-coated vesicle |
| Cellular component | GO:0030135 | 0 | 13.92 | 1 | 11 | 23 | Coated vesicle |
| Cellular component | GO:0016023 | 0 | 9.945 | 2 | 13 | 33 | Cytoplasmic membrane-bounded vesicle |
| Cellular component | GO:0031988 | 0 | 9.468 | 2 | 13 | 34 | Membrane-bounded vesicle |
| Cellular component | GO:0005886 | 0 | 2.95 | 18 | 42 | 279 | Plasma membrane |
| Cellular component | GO:0031410 | 0 | 8.276 | 2 | 13 | 37 | Cytoplasmic vesicle |
| Cellular component | GO:0031982 | 0 | 7.349 | 3 | 13 | 40 | Vesicle |
| Cellular component | GO:0034702 | 0 | 8.874 | 2 | 10 | 27 | Ion channel complex |
| Cellular component | GO:0034703 | 0 | 13.311 | 1 | 8 | 17 | Cation channel complex |
| Cellular component | GO:0015629 | 0 | 6.28 | 3 | 12 | 41 | Actin cytoskeleton |

| | | | | | | | |
|---|---|---|---|---|---|---|---|
| Cellular component | GO:0044425 | 0 | 2.116 | 34 | 59 | 532 | Membrane part |
| Cellular component | GO:0005865 | 0 | Inf | 0 | 4 | 4 | Striated muscle thin filament |
| Cellular component | GO:0031224 | 0 | 2.065 | 26 | 45 | 399 | Intrinsic to membrane |
| Cellular component | GO:0005859 | 0 | 58.893 | 0 | 4 | 5 | Muscle myosin complex |
| Cellular component | GO:0030018 | 0 | 58.893 | 0 | 4 | 5 | Z disc |
| Cellular component | GO:0031674 | 0 | 58.893 | 0 | 4 | 5 | I band |
| Cellular component | GO:0044459 | 0 | 2.58 | 11 | 25 | 176 | Plasma membrane part |
| Cellular component | GO:0016021 | 0 | 2.026 | 26 | 44 | 395 | Integral to membrane |
| Cellular component | GO:0016460 | 0 | 29.437 | 0 | 4 | 6 | Myosin II complex |
| Cellular component | GO:0048786 | 0 | Inf | 0 | 3 | 3 | Presynaptic active zone |
| Cellular component | GO:0008076 | 0 | 12.309 | 1 | 5 | 11 | Voltage-gated potassium channel complex |
| Cellular component | GO:0034705 | 0 | 12.309 | 1 | 5 | 11 | Potassium channel complex |
| Cellular component | GO:0043005 | 0.001 | 6.12 | 2 | 7 | 24 | Neuron projection |
| Cellular component | GO:0045211 | 0.001 | 6.835 | 1 | 6 | 19 | Postsynaptic membrane |
| Cellular component | GO:0005861 | 0.004 | Inf | 0 | 2 | 2 | Troponin complex |
| Cellular component | GO:0005862 | 0.004 | Inf | 0 | 2 | 2 | Muscle thin filament tropomyosin |
| Cellular component | GO:0042995 | 0.005 | 2.984 | 4 | 10 | 60 | Cell projection |
| Cellular component | GO:0005887 | 0.005 | 2.329 | 7 | 15 | 112 | Integral to plasma membrane |
| Cellular component | GO:0031226 | 0.005 | 2.329 | 7 | 15 | 112 | Intrinsic to plasma membrane |
| Cellular component | GO:0016459 | 0.008 | 6.526 | 1 | 4 | 13 | Myosin complex |
| Cellular component | GO:0030425 | 0.008 | 6.526 | 1 | 4 | 13 | Dendrite |
| Molecular function | GO:0046873 | 0 | 7.384 | 4 | 20 | 64 | Metal ion transmembrane transporter activity |
| Molecular function | GO:0022836 | 0 | 6.909 | 4 | 18 | 60 | Gated channel activity |

| | | | | | | | |
|---|---|---|---|---|---|---|---|
| Molecular function | GO:0008324 | 0 | 3.782 | 11 | 32 | 173 | Cation transmembrane transporter activity |
| Molecular function | GO:0015075 | 0 | 3.287 | 13 | 35 | 213 | Ion transmembrane transporter activity |
| Molecular function | GO:0005261 | 0 | 6.924 | 3 | 16 | 53 | Cation channel activity |
| Molecular function | GO:0005215 | 0 | 2.646 | 23 | 50 | 375 | Transporter activity |
| Molecular function | GO:0005216 | 0 | 5.473 | 5 | 19 | 75 | Ion channel activity |
| Molecular function | GO:0022857 | 0 | 2.713 | 20 | 45 | 327 | Transmembrane transporter activity |
| Molecular function | GO:0022838 | 0 | 5.19 | 5 | 19 | 78 | Substrate-specific channel activity |
| Molecular function | GO:0005509 | 0 | 4.407 | 6 | 22 | 103 | Calcium ion binding |
| Molecular function | GO:0015267 | 0 | 4.856 | 5 | 19 | 82 | Channel activity |
| Molecular function | GO:0022803 | 0 | 4.856 | 5 | 19 | 82 | Passive transmembrane transporter activity |
| Molecular function | GO:0022891 | 0 | 2.745 | 17 | 39 | 277 | Substrate-specific transmembrane transporter activity |
| Molecular function | GO:0022892 | 0 | 2.638 | 19 | 41 | 302 | Substrate-specific transporter activity |
| Molecular function | GO:0005244 | 0 | 8.671 | 2 | 11 | 31 | Voltage-gated ion channel activity |
| Molecular function | GO:0022832 | 0 | 8.671 | 2 | 11 | 31 | Voltage-gated channel activity |
| Molecular function | GO:0022843 | 0 | 9.824 | 2 | 10 | 26 | Voltage-gated cation channel activity |
| Molecular function | GO:0005515 | 0 | 1.945 | 57 | 89 | 919 | Protein binding |
| Molecular function | GO:0005267 | 0 | 11.357 | 1 | 8 | 19 | Potassium channel activity |
| Molecular function | GO:0003700 | 0 | 2.931 | 11 | 26 | 170 | Transcription factor activity |
| Molecular function | GO:0005388 | 0 | Inf | 0 | 4 | 4 | Calcium-transporting ATPase activity |
| Molecular function | GO:0015085 | 0 | Inf | 0 | 4 | 4 | Calcium ion transmembrane transporter activity |
| Molecular function | GO:0005249 | 0 | 10.892 | 1 | 7 | 17 | Voltage-gated potassium channel activity |

| | | | | | | | |
|---|---|---|---|---|---|---|---|
| Molecular function | GO:0008307 | 0 | 25.779 | 0 | 5 | 8 | Structural constituent of muscle |
| Molecular function | GO:0005184 | 0 | 7.772 | 1 | 7 | 21 | Neuropeptide hormone activity |
| Molecular function | GO:0000146 | 0 | Inf | 0 | 3 | 3 | Microfilament motor activity |
| Molecular function | GO:0015276 | 0.001 | 4.787 | 2 | 8 | 34 | Ligand-gated ion channel activity |
| Molecular function | GO:0022834 | 0.001 | 4.787 | 2 | 8 | 34 | Ligand-gated channel activity |
| Molecular function | GO:0005283 | 0.001 | 46.072 | 0 | 3 | 4 | Sodium:amino acid symporter activity |
| Molecular function | GO:0022804 | 0.001 | 2.275 | 11 | 22 | 176 | Active transmembrane transporter activity |
| Molecular function | GO:0008092 | 0.001 | 2.684 | 6 | 15 | 103 | Cytoskeletal protein binding |
| Molecular function | GO:0005488 | 0.001 | 1.504 | 161 | 184 | 2603 | Binding |
| Molecular function | GO:0005516 | 0.002 | 7.72 | 1 | 5 | 15 | Calmodulin binding |
| Molecular function | GO:0019829 | 0.002 | 7.72 | 1 | 5 | 15 | Cation-transporting ATPase activity |
| Molecular function | GO:0016820 | 0.002 | 3.245 | 4 | 11 | 64 | Hydrolase activity, acting on acid anhydrides, catalyzing transmembrane movement of substances |
| Molecular function | GO:0004723 | 0.002 | 23.03 | 0 | 3 | 5 | Calcium-dependent protein serine/threonine phosphatase activity |
| Molecular function | GO:0005217 | 0.002 | 23.03 | 0 | 3 | 5 | Intracellular ligand-gated ion channel activity |
| Molecular function | GO:0005544 | 0.002 | 23.03 | 0 | 3 | 5 | Calcium-dependent phospholipid binding |
| Molecular function | GO:0015082 | 0.002 | 10.265 | 1 | 4 | 10 | Di-, tri-valent inorganic cation transmembrane transporter activity |
| Molecular function | GO:0005179 | 0.003 | 4.341 | 2 | 7 | 32 | Hormone activity |
| Molecular function | GO:0015662 | 0.003 | 4.341 | 2 | 7 | 32 | ATPase activity, coupled to |

| | | | | | | | |
|---|---|---|---|---|---|---|---|
| | | | | | | | transmembrane movement of ions, phosphorylative mechanism |
| Molecular function | GO:0005088 | 0.003 | 6.43 | 1 | 5 | 17 | Ras guanyl-nucleotide exchange factor activity |
| Molecular function | GO:0003704 | 0.003 | 3.765 | 3 | 8 | 41 | Specific RNA polymerase II transcription factor activity |
| Molecular function | GO:0005227 | 0.004 | Inf | 0 | 2 | 2 | Calcium activated cation channel activity |
| Molecular function | GO:0005313 | 0.004 | Inf | 0 | 2 | 2 | L-glutamate transmembrane transporter activity |
| Molecular function | GO:0005523 | 0.004 | Inf | 0 | 2 | 2 | Tropomyosin binding |
| Molecular function | GO:0015172 | 0.004 | Inf | 0 | 2 | 2 | Acidic amino acid transmembrane transporter activity |
| Molecular function | GO:0015183 | 0.004 | Inf | 0 | 2 | 2 | L-aspartate transmembrane transporter activity |
| Molecular function | GO:0015269 | 0.004 | Inf | 0 | 2 | 2 | Calcium-activated potassium channel activity |
| Molecular function | GO:0015501 | 0.004 | Inf | 0 | 2 | 2 | Glutamate:sodium symporter activity |
| Molecular function | GO:0022839 | 0.004 | Inf | 0 | 2 | 2 | Ion gated channel activity |
| Molecular function | GO:0005343 | 0.004 | 15.35 | 0 | 3 | 6 | Organic acid:sodium symporter activity |
| Molecular function | GO:0042626 | 0.004 | 2.996 | 4 | 10 | 62 | ATPase activity, coupled to transmembrane movement of substances |
| Molecular function | GO:0043492 | 0.004 | 2.996 | 4 | 10 | 62 | ATPase activity, coupled to movement of substances |
| Molecular function | GO:0030528 | 0.004 | 1.762 | 20 | 32 | 323 | Transcription regulator activity |
| Molecular function | GO:0042625 | 0.005 | 3.873 | 2 | 7 | 35 | ATPase activity, coupled to transmembrane movement of ions |

| | | | | | | | |
|---|---|---|---|---|---|---|---|
| Molecular function | GO:0005543 | 0.006 | 5.14 | 1 | 5 | 20 | Phospholipid binding |
| Molecular function | GO:0015171 | 0.006 | 4.215 | 2 | 6 | 28 | Amino acid transmembrane transporter activity |
| Molecular function | GO:0043565 | 0.007 | 2.282 | 7 | 14 | 110 | Sequence-specific DNA binding |
| Molecular function | GO:0030695 | 0.008 | 2.682 | 4 | 10 | 68 | GTPase regulator activity |
| Molecular function | GO:0005089 | 0.009 | 6.153 | 1 | 4 | 14 | Rho guanyl-nucleotide exchange factor activity |
| Molecular function | GO:0005262 | 0.009 | 6.153 | 1 | 4 | 14 | Calcium channel activity |
| Molecular function | GO:0060589 | 0.009 | 2.636 | 4 | 10 | 69 | Nucleoside-triphosphatase regulator activity |
| Molecular function | GO:0042803 | 0.01 | 4.533 | 1 | 5 | 22 | Protein homodimerization activity |

Table S6: Overrepresented Gene Ontology database terms for transcripts that were down-regulated in MLX individuals relative to controls. There are a number of terms associated with, among other things, the cell cycle and chromosomal damage repair.

| Domain | ID | P-value | Odds ratio | Expected count | Observed count | Size | Term |
|---|---|---|---|---|---|---|---|
| Biological process | GO:0006259 | 0 | 5.029 | 5 | 20 | 152 | DNA metabolic process |
| Biological process | GO:0051276 | 0 | 3.875 | 6 | 19 | 179 | Chromosome organization |
| Biological process | GO:0006260 | 0 | 5.636 | 3 | 12 | 79 | DNA replication |
| Biological process | GO:0016043 | 0 | 2.24 | 29 | 50 | 877 | Cellular component organization |
| Biological process | GO:0006996 | 0 | 2.528 | 16 | 33 | 482 | Organelle organization |
| Biological process | GO:0006277 | 0 | 15.069 | 0 | 5 | 15 | DNA amplification |
| Biological process | GO:0006261 | 0 | 6.287 | 2 | 8 | 47 | DNA-dependent DNA replication |
| Biological process | GO:0006281 | 0 | 5.436 | 2 | 9 | 60 | DNA repair |
| Biological process | GO:0007049 | 0 | 2.545 | 11 | 24 | 333 | Cell cycle |
| Biological process | GO:0022402 | 0 | 2.665 | 9 | 21 | 276 | Cell cycle process |
| Biological process | GO:0044260 | 0 | 1.921 | 41 | 60 | 1230 | Cellular macromolecule metabolic process |
| Biological process | GO:0065004 | 0 | 7.899 | 1 | 6 | 29 | Protein-DNA complex assembly |
| Biological process | GO:0006974 | 0 | 4.689 | 2 | 9 | 68 | Response to DNA damage stimulus |
| Biological process | GO:0006342 | 0 | 7.568 | 1 | 6 | 30 | Chromatin silencing |
| Biological process | GO:0045814 | 0 | 7.568 | 1 | 6 | 30 | Negative regulation of gene expression, epigenetic |
| Biological process | GO:0034645 | 0 | 2.082 | 21 | 36 | 624 | Cellular macromolecule biosynthetic process |
| Biological process | GO:0009059 | 0 | 2.074 | 21 | 36 | 626 | Macromolecule biosynthetic process |
| Biological process | GO:0000278 | 0 | 2.747 | 8 | 18 | 227 | Mitotic cell cycle |
| Biological process | GO:0006270 | 0.001 | 13.29 | 0 | 4 | 13 | DNA replication initiation |
| Biological process | GO:0006323 | 0.001 | 5.318 | 2 | 7 | 47 | DNA packaging |

| | | | | | | | |
|---|---|---|---|---|---|---|---|
| Biological process | GO:0006139 | 0.001 | 1.893 | 28 | 43 | 827 | Nucleobase, nucleoside, nucleotide and nucleic acid metabolic process |
| Biological process | GO:0043170 | 0.001 | 1.783 | 49 | 67 | 1484 | Macromolecule metabolic process |
| Biological process | GO:0006807 | 0.001 | 1.84 | 31 | 47 | 938 | Nitrogen compound metabolic process |
| Biological process | GO:0009058 | 0.001 | 1.872 | 27 | 42 | 811 | Biosynthetic process |
| Biological process | GO:0006267 | 0.001 | 22.278 | 0 | 3 | 7 | Pre-replicative complex assembly |
| Biological process | GO:0071103 | 0.001 | 4.943 | 2 | 7 | 50 | DNA conformation change |
| Biological process | GO:0022403 | 0.001 | 2.492 | 8 | 18 | 247 | Cell cycle phase |
| Biological process | GO:0044249 | 0.001 | 1.877 | 26 | 41 | 787 | Cellular biosynthetic process |
| Biological process | GO:0033554 | 0.001 | 3.536 | 3 | 10 | 97 | Cellular response to stress |
| Biological process | GO:0065003 | 0.001 | 3.101 | 4 | 12 | 132 | Macromolecular complex assembly |
| Biological process | GO:0043933 | 0.002 | 2.888 | 5 | 13 | 153 | Macromolecular complex subunit organization |
| Biological process | GO:0051325 | 0.002 | 6.827 | 1 | 5 | 27 | Interphase |
| Biological process | GO:0051329 | 0.002 | 6.827 | 1 | 5 | 27 | Interphase of mitotic cell cycle |
| Biological process | GO:0007094 | 0.002 | 17.818 | 0 | 3 | 8 | Mitotic cell cycle spindle assembly checkpoint |
| Biological process | GO:0031577 | 0.002 | 17.818 | 0 | 3 | 8 | Spindle checkpoint |
| Biological process | GO:0071173 | 0.002 | 17.818 | 0 | 3 | 8 | Spindle assembly checkpoint |
| Biological process | GO:0071174 | 0.002 | 17.818 | 0 | 3 | 8 | Mitotic cell cycle spindle checkpoint |
| Biological process | GO:0009889 | 0.002 | 2.106 | 13 | 24 | 391 | Regulation of biosynthetic process |
| Biological process | GO:0031326 | 0.002 | 2.106 | 13 | 24 | 391 | Regulation of cellular biosynthetic process |
| Biological process | GO:0000075 | 0.002 | 8.532 | 1 | 4 | 18 | Cell cycle checkpoint |
| Biological process | GO:0045786 | 0.002 | 8.532 | 1 | 4 | 18 | Negative regulation of cell cycle |
| Biological process | GO:0045839 | 0.003 | 14.844 | 0 | 3 | 9 | Negative regulation of mitosis |

| | | | | | | | |
|---|---|---|---|---|---|---|---|
| Biological process | GO:0045841 | 0.003 | 14.844 | 0 | 3 | 9 | Negative regulation of mitotic metaphase/anaphase transition |
| Biological process | GO:0051784 | 0.003 | 14.844 | 0 | 3 | 9 | Negative regulation of nuclear division |
| Biological process | GO:0045449 | 0.003 | 2.127 | 11 | 21 | 335 | Regulation of transcription |
| Biological process | GO:0042990 | 0.003 | 7.961 | 1 | 4 | 19 | Regulation of transcription factor import into nucleus |
| Biological process | GO:0042991 | 0.003 | 7.961 | 1 | 4 | 19 | Transcription factor import into nucleus |
| Biological process | GO:0006271 | 0.003 | 58.984 | 0 | 2 | 3 | DNA strand elongation during DNA replication |
| Biological process | GO:0006359 | 0.003 | 58.984 | 0 | 2 | 3 | Regulation of transcription from RNA polymerase III promoter |
| Biological process | GO:0022616 | 0.003 | 58.984 | 0 | 2 | 3 | DNA strand elongation |
| Biological process | GO:0050794 | 0.004 | 1.715 | 31 | 45 | 938 | Regulation of cellular process |
| Biological process | GO:0007062 | 0.005 | 11.127 | 0 | 3 | 11 | Sister chromatid cohesion |
| Biological process | GO:0009891 | 0.005 | 3.328 | 3 | 8 | 81 | Positive regulation of biosynthetic process |
| Biological process | GO:0031328 | 0.005 | 3.328 | 3 | 8 | 81 | Positive regulation of cellular biosynthetic process |
| Biological process | GO:0042306 | 0.005 | 6.629 | 1 | 4 | 22 | Regulation of protein import into nucleus |
| Biological process | GO:0010556 | 0.005 | 1.978 | 12 | 22 | 375 | Regulation of macromolecule biosynthetic process |
| Biological process | GO:0000070 | 0.006 | 4.996 | 1 | 5 | 35 | Mitotic sister chromatid segregation |
| Biological process | GO:0016569 | 0.006 | 4.996 | 1 | 5 | 35 | Covalent chromatin modification |
| Biological process | GO:0016570 | 0.006 | 4.996 | 1 | 5 | 35 | Histone modification |
| Biological process | GO:0048477 | 0.006 | 2.262 | 7 | 15 | 221 | Oogenesis |
| Biological process | GO:0000819 | 0.006 | 4.833 | 1 | 5 | 36 | Sister chromatid segregation |
| Biological process | GO:0002781 | 0.006 | 29.484 | 0 | 2 | 4 | Antifungal peptide production |

| | | | | | | | |
|---|---|---|---|---|---|---|---|
| Biological process | GO:0002783 | 0.006 | 29.484 | 0 | 2 | 4 | Antifungal peptide biosynthetic process |
| Biological process | GO:0002788 | 0.006 | 29.484 | 0 | 2 | 4 | Regulation of antifungal peptide production |
| Biological process | GO:0002810 | 0.006 | 29.484 | 0 | 2 | 4 | Regulation of antifungal peptide biosynthetic process |
| Biological process | GO:0006611 | 0.006 | 29.484 | 0 | 2 | 4 | Protein export from nucleus |
| Biological process | GO:0006967 | 0.006 | 29.484 | 0 | 2 | 4 | Positive regulation of antifungal peptide biosynthetic process |
| Biological process | GO:0042992 | 0.006 | 29.484 | 0 | 2 | 4 | Negative regulation of transcription factor import into nucleus |
| Biological process | GO:0042994 | 0.006 | 29.484 | 0 | 2 | 4 | Cytoplasmic sequestering of transcription factor |
| Biological process | GO:0051220 | 0.006 | 29.484 | 0 | 2 | 4 | Cytoplasmic sequestering of protein |
| Biological process | GO:0033157 | 0.006 | 6.278 | 1 | 4 | 23 | Regulation of intracellular protein transport |
| Biological process | GO:0051223 | 0.006 | 6.278 | 1 | 4 | 23 | Regulation of protein transport |
| Biological process | GO:0070201 | 0.006 | 6.278 | 1 | 4 | 23 | Regulation of establishment of protein localization |
| Biological process | GO:0007093 | 0.006 | 9.888 | 0 | 3 | 12 | Mitotic cell cycle checkpoint |
| Biological process | GO:0007307 | 0.006 | 9.888 | 0 | 3 | 12 | Eggshell chorion gene amplification |
| Biological process | GO:0048523 | 0.006 | 2.186 | 8 | 16 | 244 | Negative regulation of cellular process |
| Biological process | GO:0044237 | 0.007 | 1.588 | 53 | 67 | 1587 | Cellular metabolic process |
| Biological process | GO:0031325 | 0.007 | 3.152 | 3 | 8 | 85 | Positive regulation of cellular metabolic process |
| Biological process | GO:0009893 | 0.007 | 3.11 | 3 | 8 | 86 | Positive regulation of metabolic process |
| Biological process | GO:0032386 | 0.007 | 5.963 | 1 | 4 | 24 | Regulation of intracellular transport |
| Biological process | GO:0046822 | 0.007 | 5.963 | 1 | 4 | 24 | Regulation of nucleocytoplasmic transport |

| | | | | | | | |
|---|---|---|---|---|---|---|---|
| Biological process | GO:0022607 | 0.008 | 2.048 | 10 | 18 | 293 | Cellular component assembly |
| Biological process | GO:0051726 | 0.008 | 2.853 | 3 | 9 | 105 | Regulation of cell cycle |
| Biological process | GO:0007292 | 0.008 | 2.195 | 8 | 15 | 227 | Female gamete generation |
| Biological process | GO:0007076 | 0.008 | 8.897 | 0 | 3 | 13 | Mitotic chromosome condensation |
| Biological process | GO:0030261 | 0.009 | 5.677 | 1 | 4 | 25 | Chromosome condensation |
| Biological process | GO:0032880 | 0.009 | 5.677 | 1 | 4 | 25 | Regulation of protein localization |
| Biological process | GO:0016458 | 0.009 | 3.681 | 2 | 6 | 55 | Gene silencing |
| Biological process | GO:0045892 | 0.009 | 3.25 | 2 | 7 | 72 | Negative regulation of transcription, DNA-dependent |
| Biological process | GO:0019219 | 0.009 | 1.872 | 13 | 22 | 393 | Regulation of nucleobase, nucleoside, nucleotide and nucleic acid metabolic process |
| Biological process | GO:0051171 | 0.01 | 1.866 | 13 | 22 | 394 | Regulation of nitrogen compound metabolic process |
| Biological process | GO:0048519 | 0.01 | 2.028 | 9 | 17 | 278 | Negative regulation of biological process |
| Biological process | GO:0009987 | 0.01 | 1.654 | 87 | 99 | 2603 | Cellular process |
| Biological process | GO:0044238 | 0.01 | 1.548 | 60 | 73 | 1789 | Primary metabolic process |
| Cellular component | GO:0044427 | 0 | 4.71 | 6 | 21 | 157 | Chromosomal part |
| Cellular component | GO:0005694 | 0 | 4.326 | 7 | 23 | 187 | Chromosome |
| Cellular component | GO:0005622 | 0 | 2.542 | 73 | 95 | 1998 | Intracellular |
| Cellular component | GO:0044424 | 0 | 2.343 | 66 | 88 | 1800 | Intracellular part |
| Cellular component | GO:0032993 | 0 | 10.788 | 1 | 7 | 25 | Protein-DNA complex |
| Cellular component | GO:0005634 | 0 | 2.054 | 33 | 52 | 909 | Nucleus |

| | | | | | | | |
|---|---|---|---|---|---|---|---|
| Cellular component | GO:0044428 | 0.001 | 2.236 | 15 | 28 | 406 | Nuclear part |
| Cellular component | GO:0044454 | 0.001 | 4.933 | 2 | 8 | 53 | Nuclear chromosome part |
| Cellular component | GO:0000796 | 0.001 | 26.914 | 0 | 3 | 6 | Condensin complex |
| Cellular component | GO:0000228 | 0.001 | 4.432 | 2 | 8 | 58 | Nuclear chromosome |
| Cellular component | GO:0000793 | 0.001 | 7.571 | 1 | 5 | 23 | Condensed chromosome |
| Cellular component | GO:0016585 | 0.001 | 7.571 | 1 | 5 | 23 | Chromatin remodeling complex |
| Cellular component | GO:0000785 | 0.001 | 3.852 | 3 | 9 | 74 | Chromatin |
| Cellular component | GO:0000126 | 0.001 | Inf | 0 | 2 | 2 | Transcription factor TFIIIB complex |
| Cellular component | GO:0008278 | 0.002 | 20.179 | 0 | 3 | 7 | Cohesin complex |
| Cellular component | GO:0000790 | 0.002 | 6.809 | 1 | 5 | 25 | Nuclear chromatin |
| Cellular component | GO:0043234 | 0.002 | 1.842 | 25 | 39 | 693 | Protein complex |
| Cellular component | GO:0005656 | 0.002 | 16.138 | 0 | 3 | 8 | Pre-replicative complex |
| Cellular component | GO:0043231 | 0.003 | 1.717 | 46 | 61 | 1248 | Intracellular membrane-bounded organelle |
| Cellular component | GO:0043227 | 0.003 | 1.715 | 46 | 61 | 1249 | Membrane-bounded organelle |
| Cellular component | GO:0031519 | 0.003 | 13.444 | 0 | 3 | 9 | PcG protein complex |
| Cellular component | GO:0044422 | 0.004 | 1.731 | 31 | 44 | 835 | Organelle part |

| | | | | | | | |
|---|---|---|---|---|---|---|---|
| Cellular component | GO:0044446 | 0.004 | 1.731 | 31 | 44 | 835 | Intracellular organelle part |
| Cellular component | GO:0043229 | 0.005 | 1.649 | 54 | 68 | 1465 | Intracellular organelle |
| Cellular component | GO:0043226 | 0.005 | 1.645 | 54 | 68 | 1467 | Organelle |
| Cellular component | GO:0032991 | 0.006 | 1.674 | 31 | 44 | 855 | Macromolecular complex |
| Cellular component | GO:0035098 | 0.008 | 26.692 | 0 | 2 | 4 | ESC/E(Z) complex |
| Cellular component | GO:0043228 | 0.008 | 1.81 | 17 | 27 | 463 | Non-membrane-bounded organelle |
| Cellular component | GO:0043232 | 0.008 | 1.81 | 17 | 27 | 463 | Intracellular non-membrane-bounded organelle |
| Cellular component | GO:0005654 | 0.009 | 2.391 | 6 | 12 | 152 | Nucleoplasm |
| Molecular function | GO:0008134 | 0 | 6.496 | 2 | 9 | 54 | Transcription factor binding |
| Molecular function | GO:0003677 | 0 | 2.29 | 14 | 27 | 434 | DNA binding |
| Molecular function | GO:0003682 | 0.001 | 5.208 | 2 | 7 | 50 | Chromatin binding |
| Molecular function | GO:0005542 | 0.001 | Inf | 0 | 2 | 2 | Folic acid binding |
| Molecular function | GO:0032182 | 0.002 | 5.156 | 1 | 6 | 43 | Small conjugating protein binding |
| Molecular function | GO:0032183 | 0.002 | 5.156 | 1 | 6 | 43 | SUMO binding |
| Molecular function | GO:0043138 | 0.002 | 15.654 | 0 | 3 | 9 | 3'-5' DNA helicase activity |
| Molecular function | GO:0016015 | 0.003 | 62.224 | 0 | 2 | 3 | Morphogen activity |
| Molecular function | GO:0046976 | 0.003 | 62.224 | 0 | 2 | 3 | Histone methyltransferase activity (H3-K27 specific) |
| Molecular function | GO:0005515 | 0.003 | 1.739 | 29 | 43 | 919 | Protein binding |
| Molecular function | GO:0046974 | 0.006 | 31.104 | 0 | 2 | 4 | Histone methyltransferase activity (H3-K9 specific) |

| | | | | | | | |
|---|---|---|---|---|---|---|---|
| Molecular function | GO:0008757 | 0.006 | 4.785 | 1 | 5 | 38 | S-adenosylmethionine-dependent methyltransferase activity |
| Molecular function | GO:0035251 | 0.009 | 20.731 | 0 | 2 | 5 | UDP-glucosyltransferase activity |
| Molecular function | GO:0046527 | 0.009 | 20.731 | 0 | 2 | 5 | Glucosyltransferase activity |

#### Interaction analyses

A large proportion of the significant genes participated in small interaction networks, with only two larger networks consisting of 31 and 24 genes respectively. The first network seems to be associated with muscle development and function, and the second with basic developmental functions including morphogenesis. Otherwise most interactions were linear in nature (Figure S4).

Only about half of the genes retained for analysis here (3045) overlap with those that were analysed previously (Abbott et al. 2013), indicating that these datasets have a rather different composition. However, of the genes that are in common across the datasets the correlation in estimated expression is high ($r = 0.611$, $P < 2.2*10^{-16}$). Only 5 genes were identified as significantly different in both datasets, and the direction of change was the same in all cases, and estimated magnitude was similar (Table S7). One explanation for the low number of overlapping genes could be if the transcripts identified as significant in each dataset participate in the same interaction networks, but individual significance levels vary depending on the composition of the dataset (Huang et al. 2012). However an interaction analysis implemented in NetVenn (Wang et al. 2014) suggests that this is not the case (Figure S5).

Table S7: Genes that were identified as having changed significantly in this dataset (MLX) and a previously analysed one (Abbott et al. 2013). The direction and magnitude of change is generally consistent.

| Probe ID | Gene name | Change in imprinting dataset | Change in MLX dataset |
|---|---|---|---|
| **1632656_at** | **CG12112** | -1.479302589 | -1.0523 |
| **1626392_s_at** | **Cyp4aa1** | 1.085289793 | 0.871412 |
| **1627068_at** | **CG8319** | 0.661599145 | 0.661458 |
| **1635183_at** | **Mef2** | -1.14961547 | -0.63993 |
| **1638886_at** | **Spn43Ab** | -0.74362116 | -0.53079 |

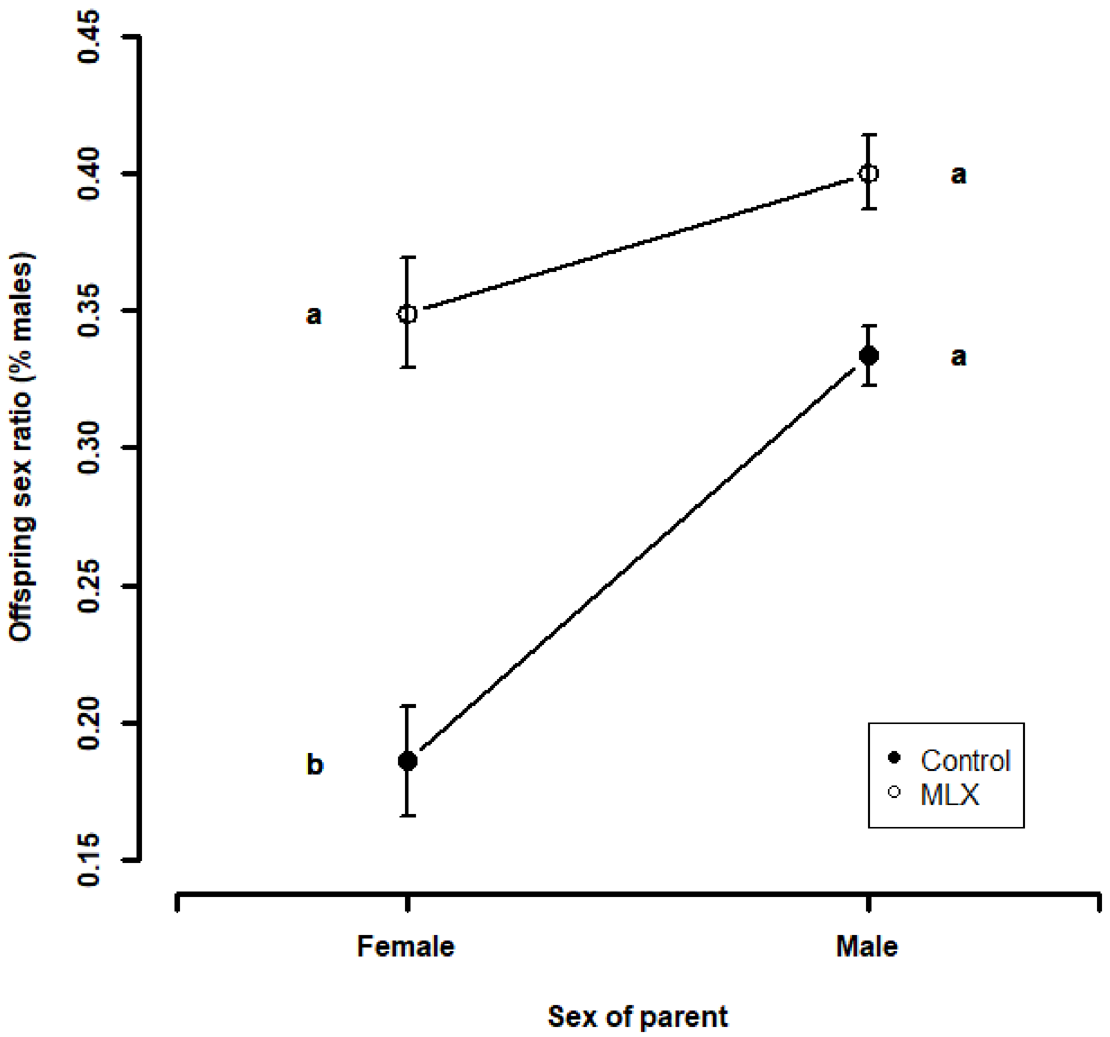


Figure S1: Sex ratio in relation to experimental evolution treatment and parental sex. Plot shows means and standard errors. Homogenous subsets are marked with “a” and “b”. Individuals carrying an MLX X-chromosome produce a higher proportion of male offspring. Homogenous subsets are indicated with letters. The pairwise comparison between Control and MLX males was nearly significant ($P$ = 0.0544), consistent with a main effect of the selection treatment. Note that the lower values for females are because the data was collected two days later, and some male mortality was observed to have occurred during that period.

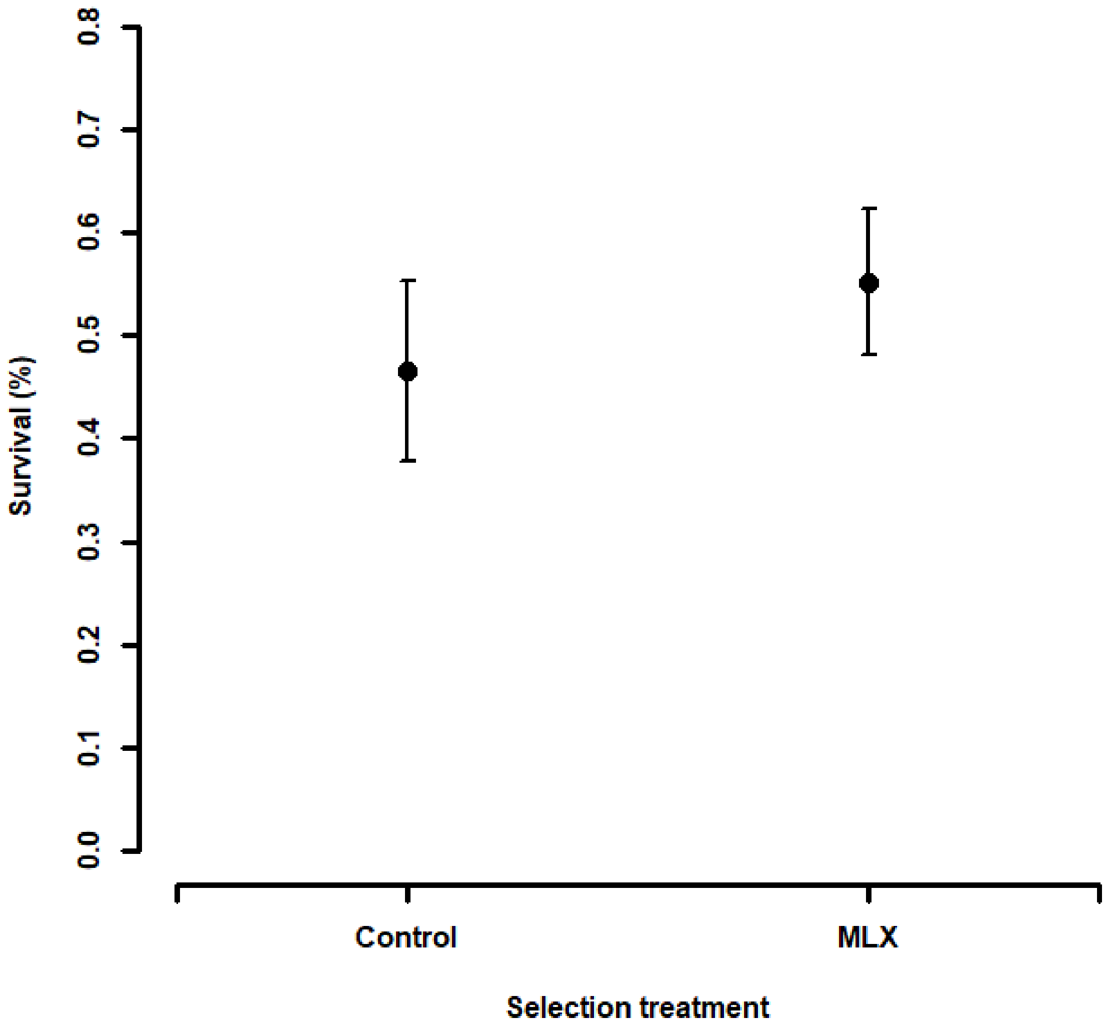


Figure S2: Difference in offspring survival according to selection treatment. Plot shows means and standard errors. Offspring of MLX females have significantly higher egg-to-adult survival.

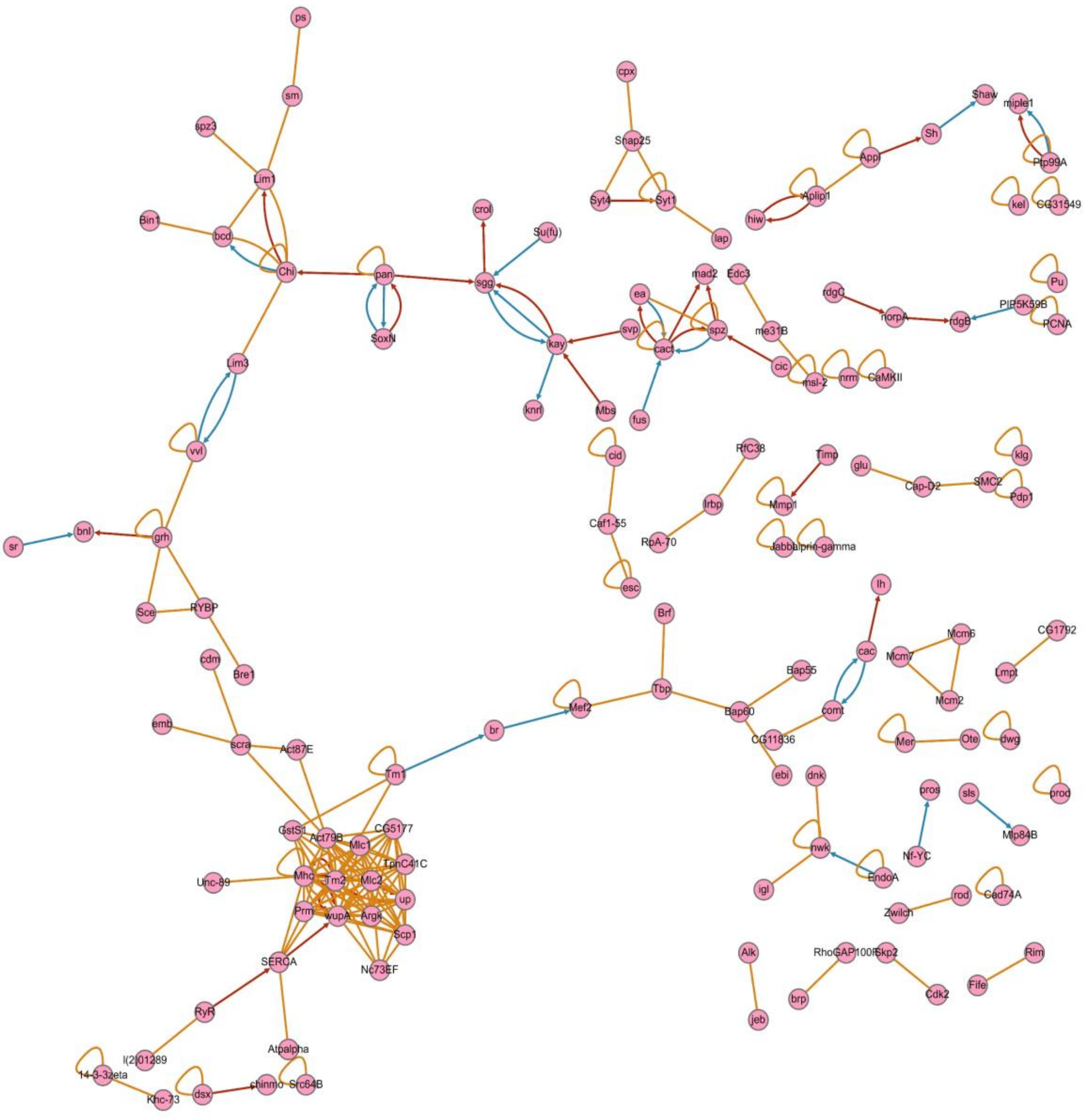


Figure S3: Network analysis of interactions between genes identified as significantly different in this dataset. There is one large network associated with muscle development and function, and one associated with basic developmental functions including morphogenesis.

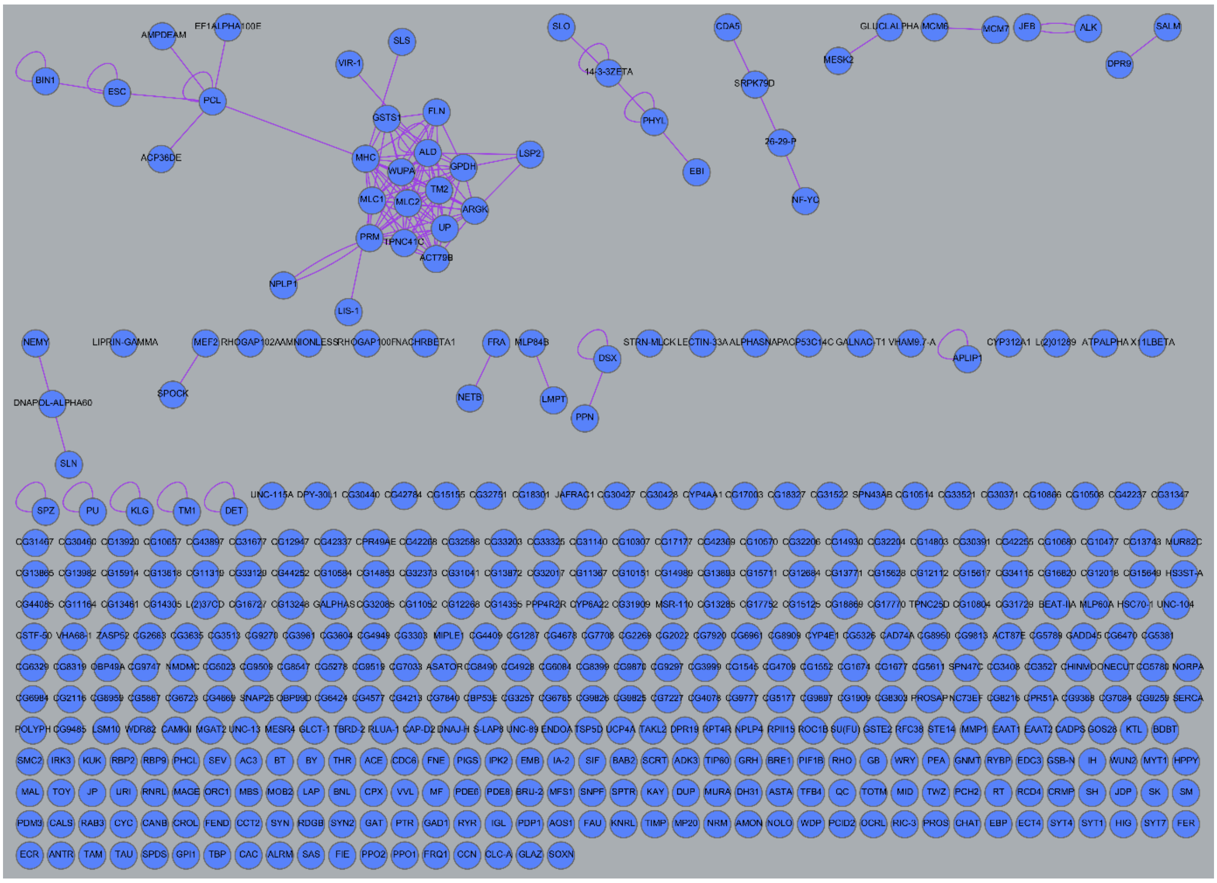

Figure S4: Network analysis of interactions between genes identified as significantly different in this dataset, and in a partially overlapping previous one (Abbott et al. 2013). It is clear that the majority of genes do not interact with each other.
